# Experimentally-induced and real-world acute anxiety have no effect on goal-directed behaviour

**DOI:** 10.1101/606145

**Authors:** CM Gillan, MM Vaghi, FH Hezemans, Grothe S van Ghesel, J Dafflon, AB Brühl, G Savulich, TW Robbins

## Abstract

Compulsivity is associated with failures in goal-directed control, an important cognitive faculty that protects against developing habits. But might this effect be explained by co-occurring anxiety? Previous studies have found goal-directed deficits in other anxiety disorders, and to some extent when healthy individuals are stressed, suggesting this is plausible. We carried out a causal test of this hypothesis in two experiments (between-subject N=88; within-subject N=50) that used the inhalation of hypercapnic gas (7.5% CO_2)_ to induce an acute state of anxiety in healthy volunteers. In both experiments, we successfully induced anxiety, assessed physiologically and psychologically, but this did not affect goal-directed performance. In a third experiment (N=1413), we used a correlational design to test if real-life anxiety-provoking events (panic attacks, stressful events) impair goal-directed control. While small effects were observed, none survived controlling for individual differences in compulsivity. These data suggest that anxiety has no meaningful impact on goal-directed control.

## Introduction

Two well-established systems contribute to everyday decision making and behaviour, the goal-directed and the habitual system (1). Goal-directed behaviour is characterized by actions that are appropriate to the current desire for a given outcome and informed by the knowledge of the causal relationship between an action and the associated outcome (2). More recently goal-directed control has been formalized as model-based planning, within a reinforcement learning framework (3). A fragile goal-directed system leads one to get stuck in habits (4) and typifies not only Obsessive-Compulsive Disorder (OCD) (5-7) but also several other psychiatric conditions on the compulsivity spectrum such as eating disorder, drug abuse and alcohol addiction (8, 9). Accordingly, it has been suggested that goal-directed deficits constitute a trans-diagnostic trait (10, 11).

One potential issue with this model is its specificity. Compulsivity is highly comorbid with anxiety (12), which is unsurprising, as OCD has only recently moved out of the Diagnostic and Statistical Manual category of anxiety disorders to its own classification (13). Accordingly, this raises the possibility that elevated anxiety levels in OCD might account for failures in goal-directed planning and consequent overreliance on habits. Indeed, physiological and psychological stress has been likened to anxiety, and is generally thought to impair several forms of deliberative and reflective processes, in favour of more automatic and reflexive ones (14). In support of this idea, social anxiety patients appear to show similar deficits in goal-directed planning to OCD patients, despite the fact that they do not have a compulsive phenotype (15). Cross-sectional, correlational work has started to address this issue, finding that when a range of psychopathology measures are taken (and controlled for) within the same individuals, there is no meaningful contribution of trait anxiety to goal-directed deficits, while the association with compulsivity is robust (10, 11). However, these studies are limited not just by their correlational nature, but because they assess trait anxiety, which does not speak to acute states of anxiety that are experienced by patients more transiently, often in response to their own symptoms (16).

Though no previous study has examined whether experimentally induced state anxiety impairs goal-directed planning, a related literature on stress-induction offers a basis for this suggestion. Specifically, acute stress has been shown to induce deficits in goal-directed planning (17-19), albeit inconsistently (null results: 20, 21, 22) in healthy individuals. Acute anxiety and stress manipulations produce similar cardiovascular changes, and induce negative affect, but anxiety induction differs from stress in terms of the specific psychological experience (e.g. increased vigilance, panic, fear) and other aspects of the physiological response (23, 24). The present study used a combination of causal and correlational approaches to investigate the role of acute anxiety on goal-directed control, in three experiments spanning laboratory and real-life settings.

First, we used hypercapnic gas (i.e. with increased CO_2_ level) to experimentally induce state anxiety and test its impact on goal-directed control, operationalized as sensitivity to contingency degradation (7). Hypercapnic gas is a well-validated method for experimentally inducing a transitory state of acute anxiety in healthy volunteers (25). At very high doses (35% CO_2)_ it generates symptoms similar to those of panic disorder, with increased blood pressure and bradycardia (26-28), especially in subjects with panic disorder or susceptibility to it (29, 30). We used lower doses (7.5% CO_2)_ which are reported to be sufficient to induce physiological and psychological symptoms of anxiety and sustained arousal associated with an anxiety state (24). Subjects had profound physiological and subjective psychological responses to the anxiety induction procedure including changes in heart rate, blood pressure and self-reported anxiety, but it failed to induce deficits in goal-directed control over behaviour.

Reasoning this might be associated with task and study design sensitivity, we repeated this experiment using a within-subjects design and a different measure of goal-directed control – a ‘model-based planning’ measure derived from the two-step reinforcement learning task described above (3). Again, the procedure had substantial physiological and psychological effects consistent with the induction of an acute state of anxiety, but this had no demonstrable detrimental effect on goal-directed behaviour.

In a third and final experiment, we tested this hypothesis in a naturalistic, real-world setting using a large-scale correlational design (N=1413) (10). We investigated if goal-directed (model-based) control is impaired in individuals who suffered recent ‘real life’ acute anxiety, specifically that known to be associated with the experience of a recent panic attack (31) and/or major life-stressors (32). We found that the recent occurrence of a panic attack and higher levels of stress in the past year were both modestly associated with deficits in goal-directed planning. Crucially, neither survived controlling for a correlated psychiatric trait, compulsive behaviour and intrusive thought, which we previously showed has a strong association with goal-directed planning using these data (10).

## Results

### Anxiety induction and Contingency Degradation (Experiment 1)

In a between-subjects design, one group was assigned to inhale hypercapnic gas (7.5% CO_2)_ during the performance of a cognitive task, while the other inhaled normal air. Psychological and physiological measures confirmed that anxiety induction was successful and of a magnitude similar to that observed in prior studies (33-35): participants in the CO_2_ condition experienced greater self-reported anxiety (*p* < .001) and had a higher heart rate (*p* < .001) than those assigned to the Air condition (Figure 1B and 1D; supplementary materials). We tested if this manipulation would affect subjects’ ability to detect action-outcome instrumental contingency using a contingency degradation paradigm (7), one of the earliest operationalisations of goal-directed learning from the animal literature (36).

**Figure 1.**
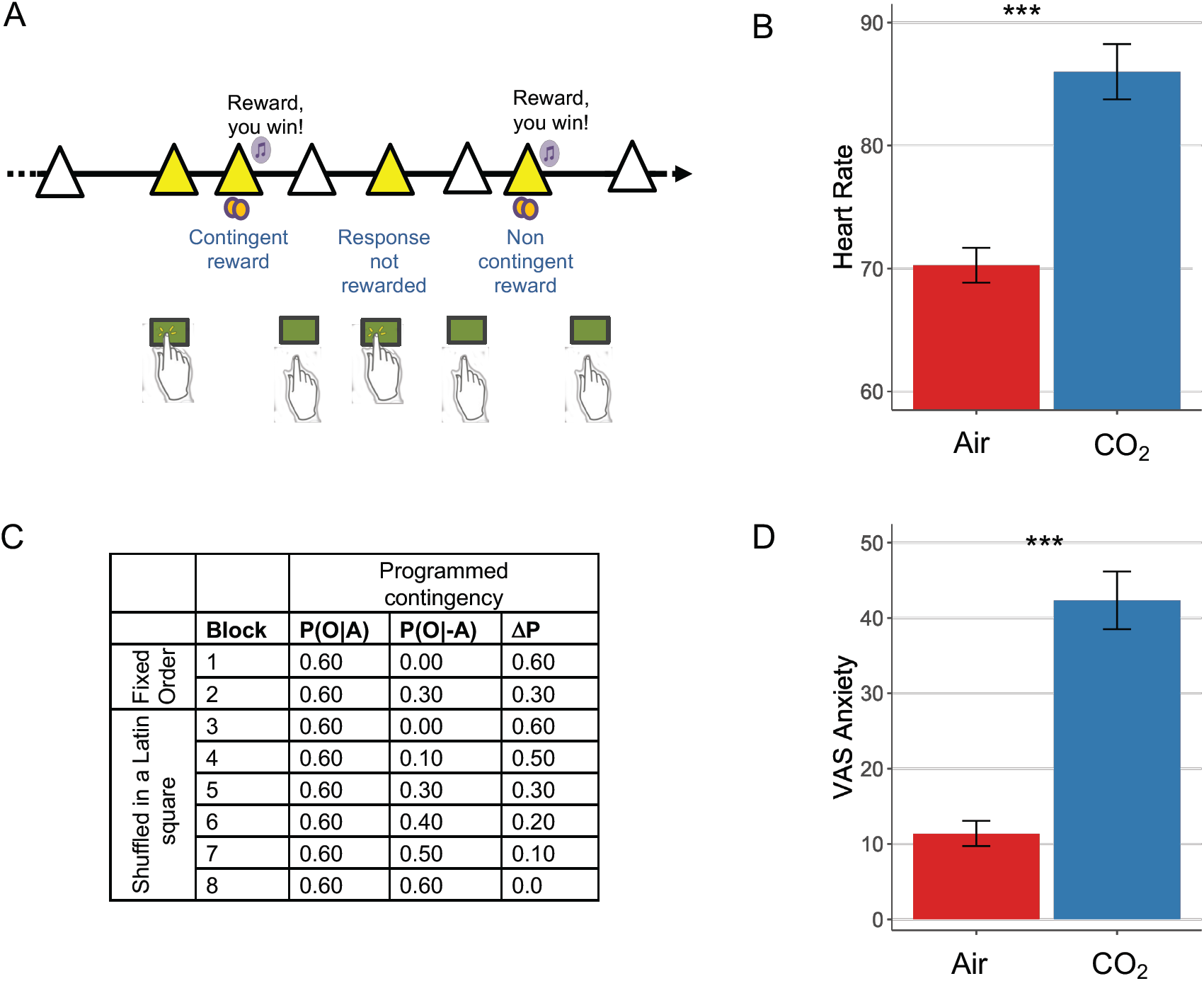
Experiment 1 Study Design. A. Contingency degradation task design. In each block, subjects were presented with a white triangle, signalling that they had the opportunity to press or to not press the space bar, in a free-operant, self-paced procedure(7). The triangle turned yellow when a response was recorded. Rewards (a 25 pence image) were delivered according to a probability, P(O|A), on trials when a response was made, and P(O|-A) when a response was not made. B. Physiological response to anxiety induction. Heart rate was elevated significantly during the gas condition, *p* < .001. C. Programmed contingencies. Each participant completed 8 blocks where contingency was systematically varied through changes to P(O|-A). The first two blocks were considered training blocks and appeared in a fixed order as denoted in the table. The 6 remaining test blocks were presented in a counterbalanced order across subjects. D. Psychological response to anxiety induction. Anxiety scores measures using a visual analogue scale (VAS) were also significantly elevated during the inhalation of gas compared with air, *p* < .001.

Participants learnt the contingencies in the training phase (*F*_(1, 86)_ = 26.48, *p* < .001). Experimentally-induced anxiety did not affect subjects’ behavioural sensitivity to instrumental contingency. Participants overall adjusted their response rate in line with the underlying contingency, as evidenced by a main effect of contingency on response rate in the test blocks (*F*_(3.73, 320.59)_ = 29.95, *p* < .001). In the test blocks, there was no between-group difference (*F*_(1, 86)_ = 0.22, *p* = .64) and no group by contingency interaction (*F*_(3.73, 320.59)_ = 1.74, *p* = .15) (Figure 2A). Bayes Factor analysis further confirmed these findings (Supplementary Material).

**Figure 2.**
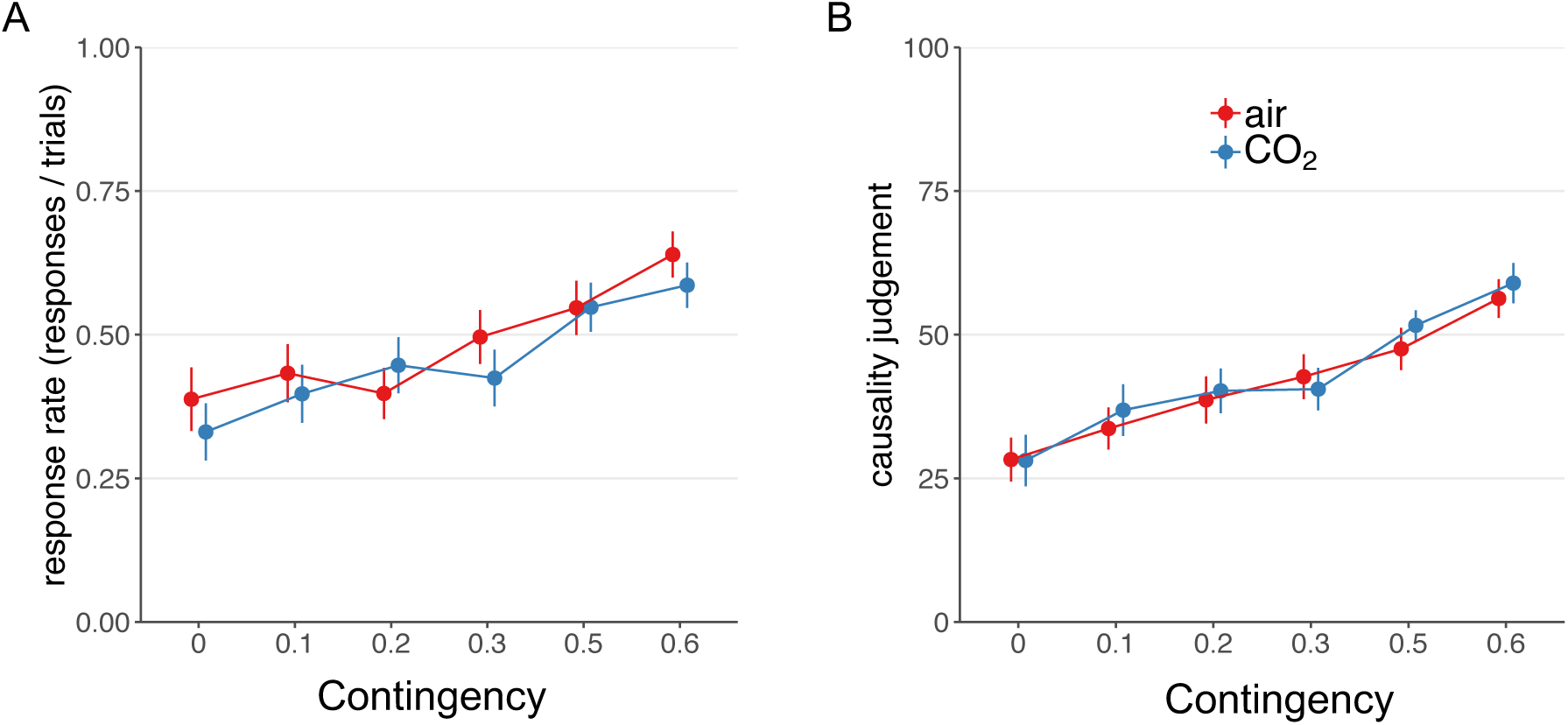
Results from Experiment 1. A. There was no effect of CO_2_-induced anxiety on subjects’ sensitivity to instrumental contingency as measured by choice responses, *F*_(3.73, 320.59)_ = 1.74, *p* = .15. B. There was similarly no effect of group on the extent to which causality judgements scaled with instrumental contingency, *F*_(2.99, 256.89)_ = 0.33, *p* = .81.

The same was true of participants’ *subjective* assessments of instrumental contingency (i.e. their explicit model of the environment). Subjects accurately tracked the underlying contingency of the task (training blocks, *F*_(1, 86)_ = 30.46, *p* < .001; test blocks, *F*_(2.99, 256.89)_ = 26.22, *p* < .001) and the experimental manipulation did not affect this. There was no between-group difference (*F*_(1, 86)_ = 0.16, *p* = .69) and no group by contingency interaction (*F*_(2.99, 256.89)_ = 0.33, *p* = .81) (Figure 2B) on causality judgements. Mirroring the findings on choice responses, experimentally-induced anxiety did not affect subjective judgments of instrumental contingency – adding weight to the suggestion that state anxiety may not have an appreciable effect on goal-directed control over action.

### Individual Differences

Prior work showed that individual differences might be important in revealing the effect of stress on goal-directed behaviour (18, 20-22). Therefore, we tested if perceived anxiety in response to CO_2_ manipulation played a role in susceptibility to detecting instrumental contingency. We ran the model explained above with programmed contingency as a within-subject factor, introducing the change in acute anxiety as a between-subject covariate. The change in acute anxiety factor was computed as the difference between VAS-anxious before inhaling the gas and after inhaling the gas. As above, there was a significant effect of programmed contingency (*F*_(3.73, 309.85)_ = 25.42, *p*< .001) on response rate, but there was no main effect of individual sensitivity (*F*_(1, 83)_=0.28, *p*=.60) to the manipulation nor an interaction effect (*F*_(3.73, 309.85)_ =0.20, *p*=.42). Similar findings were obtained on subjective causality ratings. Accordingly, programmed contingency (*F*_(3.18, 264.24)_=33.10, *p*<.001) significantly predicted causality ratings, but there was not a main effect (*F*_(1, 83)_=0.00, *p* =.96) nor a significant interaction with individual sensitivity to the manipulation (*F*_(3.18, 264.24)_=0.20, *p* =.90). Therefore, individual differences in experienced anxiety as reported by subjects upon CO_2_ challenge did not affect goal-directed planning.

### Anxiety Induction and Model-Based Planning (Experiment 2)

Experiment 1 found that experimentally-induced anxiety had no effect on the extent to which actions were flexibly updated in accordance with changes in instrumental contingency. There was similarly no effect on subjects’ ability to generate an explicit model of contingency in the environment. In experiment 2, we tested if anxiety induction would affect goal-directed planning on a complementary ‘model-based’ learning task (Figure 3) (3, 37). Model-based planning has been previously shown to correlate with sensitivity to outcome devaluation (4), OCD diagnosis (8), symptoms (10), and has been successfully modified using pharmacological manipulations (38, 39). As such, it represents a somewhat more established test of goal-directed planning. In contrast to Experiment 1, we employed a within-subjects design, which overcomes the potential problem that individual differences in goal-directed control (e.g. associated with compulsiveness, IQ, age (10)) may have hindered our ability to detect changes resulting from anxiety-induction in Experiment 1.

**Figure 3.**
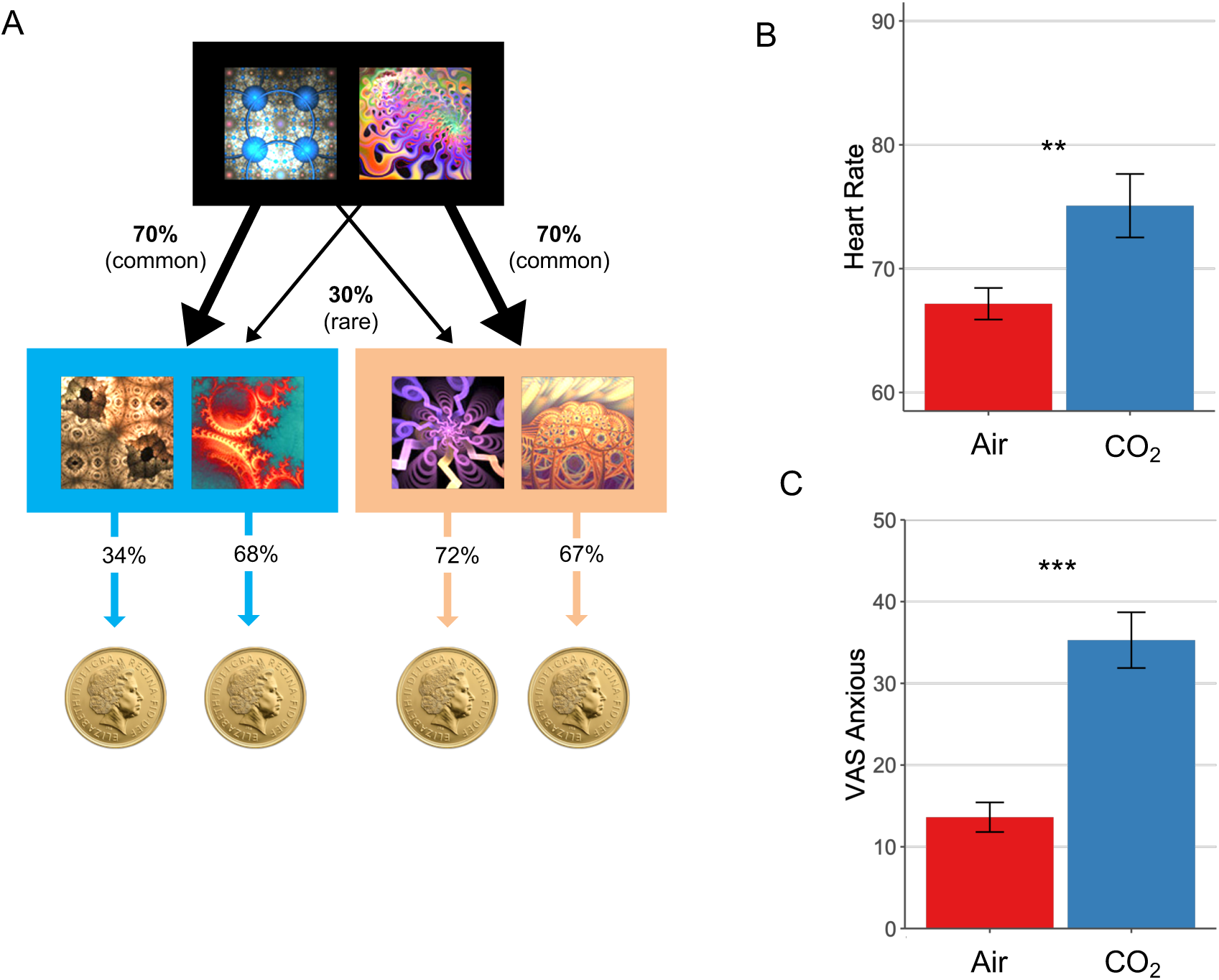
Model-Based Learning Task. A. On each trial, subjects chose between two fractals, which probabilistically transition to either an orange or blue state where they must make another choice. In this schematic, the fractal on the left had a 70% chance of transitioning to the blue state, what is called a ‘common’ transition, and a 30% chance of transitioning to the orange state, i.e. a ‘rare’ transition. In the second orange or blue state, subjects again chose between two fractals, each of which was associated with a probability of reward (a pound coin). Unlike the transition structure, these reward probabilities drifted slowly over time (.25 < P< .75). This meant that subjects were required to dynamically track which of the fractals in the orange and blue states were currently best. The reward probabilities depicted (34%, 68%, 72%, 67%) refer to an example trial. Model-based planning on this task is operationalised as the extent to which subjects’ decision to repeat an action at the first stage, depend on (i) whether this action was rewarded on the previous trial and (ii) and whether the path from action to outcome was expected (‘common’). B. Physiological response to anxiety induction. Heart rate was elevated significantly during the gas condition, *F*_(1,49)_=10.72, *p*=.002. C. Psychological response to anxiety induction. Self-reported anxiety levels were also significantly elevated during the inhalation of gas compared with air, *F*_*(1,49*)=_57.47, *p*<.001.

As in Experiment 1, the CO_2_ manipulation was effective in inducing anxiety in subjects (Figure 3B and Figure 3C), but once again, this did not alter goal-directed performance as CO_2_ had no effect on model-based planning (β=-0.03, SE=0.04, *p*=.44). The regression model overall fit subjects’ behaviour as expected; ‘model-free’ behaviour was evident in the sample (β =.55, SE=.08, *p*<.001) which refers to how much subjects tend to repeat actions that were recently rewarded. Model-based learning was also overall significant (β=.28, SE=.06, *p*<.001), such that subjects took environmental contingency into account when deciding whether or not to repeat a rewarded choice. Finally, subjects showed an overall biased tendency to repeat choices from one trial to the next, regardless of reward or transition information (β=1.59, SE=.12, *p*<.001). Much like model-based learning, there was no effect of anxiety on model-free learning (β=- 0.02, SE=0.03, *p*=.52), or action repetition (β=-0.08, SE=0.04, *p*=.060; Figure 4, Supplementary Table S5). Although the latter approached significance, such that subjects had a slight tendency to switch choices more while under CO_2_. These analyses were complemented with a full computational model, details of which are available in the online supplement, with the only difference being that the effect of CO_2_ on choice switching was significant in this more comprehensive computational analysis (Supplementary Table S8). Thus, it appears there may be a modest association between acute anxiety and an increased tendency to explore new options from trial to trial.

**Figure 4.**
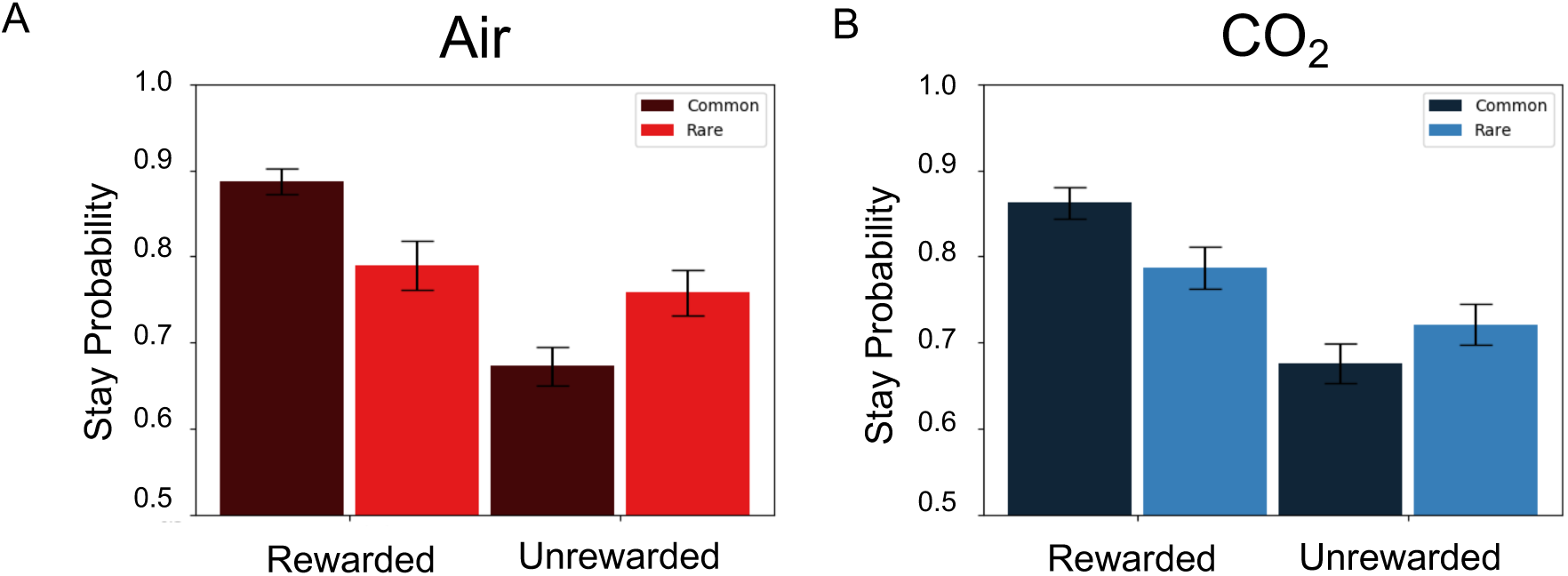
Figure 2. Results from Experiment 2. A. Stay/switch behaviour for subjects in the Air condition as a function of whether or not the last trial was rewarded/unrewarded and followed a rare/common transition. B. The same plot, showing the group average behaviour under CO_2_. In both plots, subjects showed the classic signatures of both model-based and model-free planning, indexed by a significant reward × transition interaction (β=.28, SE=.06, *p*<.001) and a main effect of reward (β =.55, SE=.08, *p*<.001).

### Individual Differences

Following the same logic as Experiment 1 – that individual differences might be important in revealing the effect of stress on goal-directed behaviour (18, 20, 21) – we tested if the effects of CO_2_ on behaviour might be detectible when we take into account how strongly subjects reacted to the CO_2_ manipulation. To do this, we entered subjects’ self-reported anxiety scores into the analysis described above as an interacting variable. As above, model-based planning was not affected by the gas/air manipulation (β=-0.03, SE=0.04, *p*=.38), and this effect did not depend on individual sensitivity to the manipulation (reward by transition by CO_2_ by anxiety change, β=-0.06, SE=0.04, *p*=.07). Although the trend was in the expected direction, this effect was not specific – those individuals who were most anxious under CO_2_ also tended to switch more under CO_2_ (β=-0.14, SE=0.04, *p*<.001).

### Real life acute anxiety (Experiment 3)

Consistent with the findings from Experiment 1, we found no effect of an acute anxiety induction on goal-directed planning in Experiment 2. In a final experiment, we tested if acute anxiety in a real-life, more ecologically valid, setting might be necessary to reveal the hypothesised detrimental effect of anxiety on goal-directed behaviour. We tested 1413 subjects online using Amazon’s Mechanical Turk on the model-based learning task described above. Findings relating to the association between compulsivity and model-based planning has been published elsewhere (10), but in data not previously published, we enquired about whether subjects had a panic attack in the past week, which is known to induce a temporary state of acute anxiety. We chose to examine panic attacks, rather than using a questionnaire probing state anxiety, because state anxiety has an unacceptably high correlation with trait anxiety when measured in the absence of an acute stressor (e.g. r=.71 (40)). As our prior work has demonstrated that trait anxiety is not related to goal-directed planning (10), we wanted to ensure that our measure of acute anxiety was not in large part confounded by trait anxiety. Measuring the occurrence of recent panic attacks is an attractive alternative (although not without limitation), because they represent an acute anxiety provoking event (31) and as such is more comparable to our lab-based anxiety induction. Criteria for a panic attack were from item 1 of a validated instrument (Panic Disorder Severity Scale, PDSS (41)) and in brief required subjects to have experienced 4 of 17 symptoms (e.g. rapid or pounding heartbeat, feeling of choking, nausea, chills or hot flushes, fear of dying) and that the panic attack must have been a “sudden rush of fear or discomfort”, peaking within 10 minutes. Episodes like panic attacks that have fewer than 4 symptoms were defined as limited symptom attacks, but also contributed to subjects’ score. Specifically, subjects indicated the frequency of panic or limited symptom attacks in the past week on item 1 of the PDSS and this served as our measure for subsequent analyses.

Consistent with other general population samples (42), approximately a third (N=474) of our online sample indicated they had experienced a panic or limited symptom attack in the past week (Figure 5A). The frequency of panic attacks in the past week was correlated with reductions in model-based planning (β=-.03, SE =.01, *p*=.012), such that individuals who had more panic attacks in the past week showed the lowest scores of model-based planning. This lends support to the notion that recent episodes of acute anxiety, here induced through panic attacks, can impair goal-directed planning. However, the occurrence of panic attacks is elevated in mental health disorders in general (43, 44), which raises the possibility that other aspects of psychopathology might account for this result. We therefore tested if compulsivity, which has been previously linked to goal-directed deficits, might explain the relationship between panic attacks and reductions in model-based planning. This was indeed the case – the effect of having a recent panic attack did not survive controlling for individual differences in “Compulsive Behaviour and Intrusive Thought”, a trans-diagnostic dimension that is reliably associated with goal-directed failures (β=-.04, SE=.01, *p*<.001; note this finding was previously published(10)). As expected, scores on the compulsive dimension were correlated with frequency of panic attacks (r=.42, *p*<.001). The effect of panic attacks on model-based planning was reduced to β=-.01, SE=.01, *p*=.33 when the compulsive factor was present in the model (Figure 5C). When we used the more elaborated computational model, the effect of panic attacks on goal-directed planning approached zero in the opposite direction (β=.003, SE=.01, *p*=.81) after compulsivity was controlled for (Supplementary Table S9).

**Figure 5.**
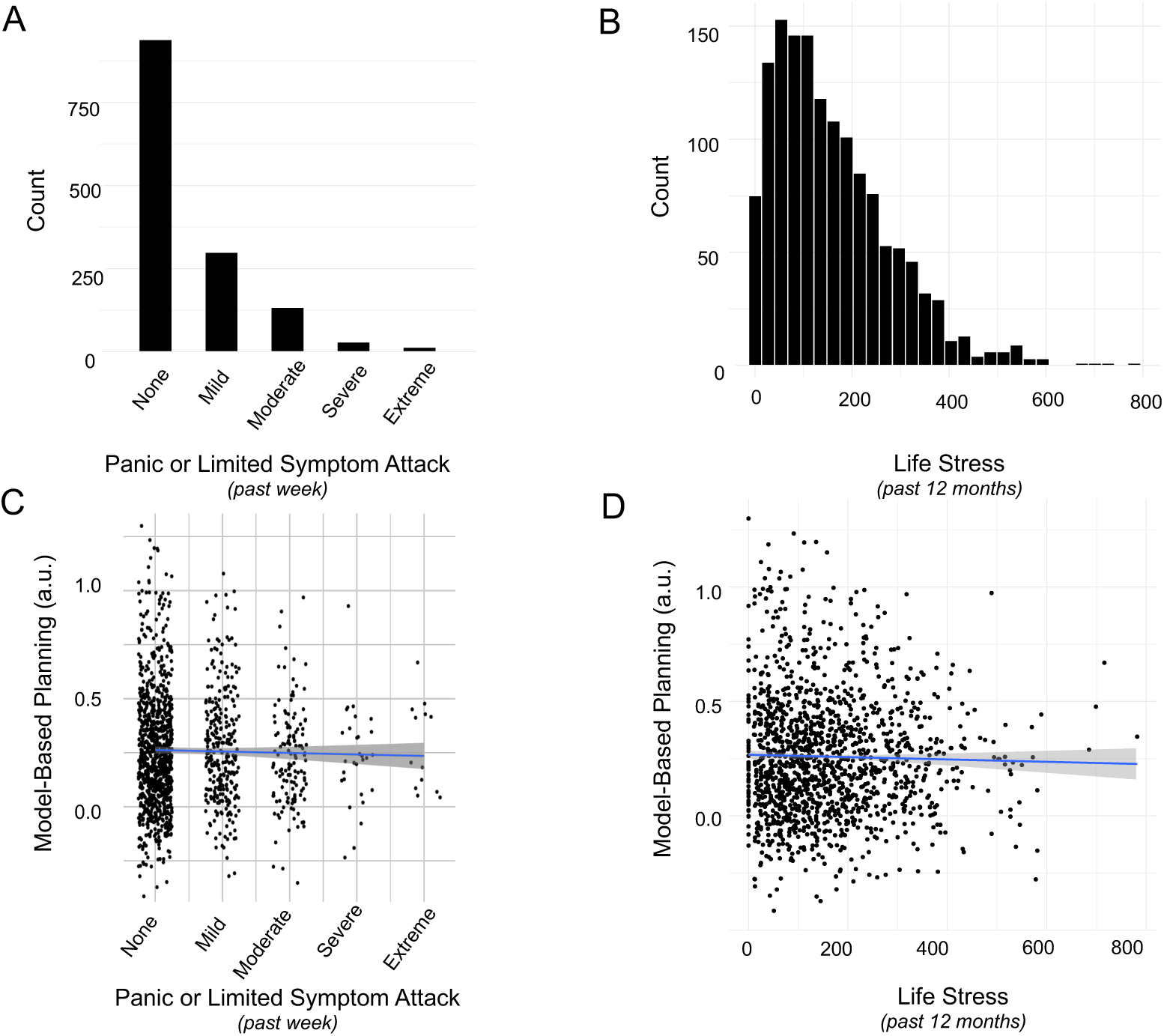
Results from Experiment 3. A. Histogram displaying the number of individuals endorsing the various levels of frequency and severity of panic attacks in the past week. Scores were coded as follows: none (“no panic or limited symptom attacks”), mild (no full panic attacks and no more than 1 limited symptom attack/day), moderate (“1 or 2 full panic attacks and/or multiple limited symptom attacks/day”), severe (Severe: more than 2 full attacks but not more than 1/day on average) and extreme (“full panic attacks occurred more than once a day, more days than not”). B. Histogram displaying the distribution of life stress scores in the sample. C. There was no association between model-based planning and the occurrence of panic attacks in the past week, after controlling for age, gender, IQ and compulsive symptomatology, β=-.01, SE=.01, *p*=.33. Y-axis displays residuals for model-based planning after these features are taken into account. D. There was no association between model-based planning and life stress experienced over the past year, after controlling for age, gender, IQ and compulsive symptomatology, β=-.01, SE=.01, *p*=.33. As above, Y-axis displays residuals for model-based planning.

We observed an association between frequency of panic attacks and choice switching (*p*=.012), mirroring our causal result from Experiment 2. However, the effect of panic attacks on increased switching did not survive inclusion of compulsivity in the model for the one-trial-back regression (*p*=.23), or in the computational model (Supplementary Table S9; *p*=.06).

Finally, we tested if life stress in the past year was associated with deficits in model-based planning. This was assessed using the Social Readjustment Scale (45), which presents an inventory of common stressful life events to participants and asks them to select those that applied to them in the previous 12 months (e.g. death of a spouse, divorce) (Figure 5B). Much like a recent panic attack, we found that life stress scores were linked to failures in model-based planning (β=-.02, SE=.01, *p*=.04). However, as was the case for panic attacks, life stress was also correlated with the compulsive factor (r=.29, *p*<.001), and indeed the relationship to model-based planning did not survive inclusion of the compulsive factor in the analysis. Specifically, the effect of life stress on model-based planning was reduced to β=-.01(SE=.01, *p*=.33; Figure 5D) in the regression analysis and β=-.01, SE=.01, *p*=.24 in the full computational model (Supplementary Table S10).

## Discussion

Across three independent experiments, we found no evidence that acute anxiety has a detrimental effect on goal-directed planning. The first two studies employed an extensively validated causal manipulation for inducing an acute state of anxiety, inhalation of air enriched with CO_2_ (24, 28). Using both between- and within-subject designs, and two well-validated and widely used tests for goal-directed behaviour, neither study found evidence in support of the hypothesis. A third study took a correlational, but larger scale (N=1413), approach and tested if individuals who recently experienced a panic attack, which is associated with an increase in acute state anxiety (31), had poorer goal-directed performance. Unlike most clinical studies, this design incorporated a comprehensive range of clinical assessments and could thus control for clinical confounds (like trait compulsivity). While we found that those who experienced more panic attacks in the past week had greater deficits in goal-directed planning, this did not survive controlling for compulsivity, which has been extensively studied in the content of goal-directed control failures. Together, these data contribute to a larger literature suggest that trait (10), and now state, anxiety do not have demonstrably detrimental effects on goal-directed planning.

The most consistent cognitive changes that have been linked to trait anxiety are an increased attentional bias to threat or ‘hypervigilance’ (46) and the tendency to interpret ambiguous stimuli as threatening (47). Results from studies using the 7.5% CO_2_ challenge closely mirror these findings – with the manipulation increasing alerting and orienting (33), threat processing (e.g. hypervigilance) (35) and negative interpretations of neutral events (34), thus suggesting that 7.5% hypercapnic gas manipulation in the lab can mirror cognitive changes observed in association with anxiety. While the putative role that anxiety plays in more complex forms of decision-making is of broad interest (48), there is a dearth of evidence suggesting it has effects that are not explained as knock-on effects of increases in threat-sensitivity and vigilance. For example, while there is some evidence to suggest that clinically anxious individuals tend to make better long-term choices e.g. on the Iowa Gambling Task (IGT), this appears to result from a bias to avoid losses, which in the context of this task is confounded with the choice of ‘advantageous’ decks (49). Even this, however, has been inconsistently shown, with another study finding that high trait anxiety leads to *impaired* IGT performance (50). One potential explanation for inconsistent results in this area is that studies have been largely cross-sectional and correlational – something the design of the present investigation overcame.

Although no previous studies have examined the effect of acute experimentally-induced state anxiety on goal-directed control, several studies studied the impact of stress (17, 18, 20-22, 51) in healthy volunteers. Three studies found that stress induced goal-directed deficits (17-19), mirroring findings in rodents following 21 days of unpredictable stress exposure (51). Three other studies, however, found no such effect (20-22). One key point of departure between studies that did and did not see an effect was type of stressor used. Those that found significant effects used a socially-evaluated cold pressor test, and those that did not used either the cold pressor in isolation (20), or a social stress test in isolation (21, 22). This distinction is important as the socially evaluated cold pressor test has been shown to induce a much stronger increase in cortisol, compared to cold pressor test alone (52), with the procedures otherwise eliciting similar cardiovascular and subjective stress responses. The notion that cortisol might mediate stress effects on goal-directed planning is supported by the observation that changes in cortisol were linked to deficits in performance in studies that failed to otherwise show a main effect of stress (20, 21). In other words, the largest increases in cortisol were linked to the largest task deficits. This ties in with pharmacological evidence showing that decrements in goal-directed performance cannot be induced through noradrenergic manipulation only; concurrent glucocorticoid stimulation is necessary (although not sufficient) (53, 54). Differential involvement of cortisol might explain why acute stress appears to have an impact on goal-directed planning, but anxiety induction does not. While acute stress and anxiety induction result in similar cardiovascular effects (i.e. increases in heart rate and blood pressure) (24, 52) and noradrenergic activation (55, 56), anxiety induction via 7.5% CO_2_ does not result in a reliable increase in cortisol (25, 57). Hypercapnia causes more pronounced and specific increases in self-reported feelings of anxiousness, fear, panic and worry, which are reduced in response to common treatments for generalized anxiety, including anxiolytics (58, 59).

The extent to which more chronic forms of real-life stress impair goal-directed control is an open question and has only been partially addressed in one prior study with a relatively small sample (N=39)(21). Subjects with high self-reported chronic stress levels had a larger effect of acute stress on model-based planning performance, than their low stress counterparts (21). This might suggest that goal-directed learning is in some sense more fragile in individuals who have high levels of chronic life stress, but this is difficult to assess as the authors did not report any test for the direct association between life stress and model-based planning. We tested this using a large sample (N=1413) and did not find evidence for an association, after controlling for compulsivity. This suggests that the impact of real-life chronic stress on goal-directed planning, if it exists, is certainly less pronounced than folk wisdom suggests.

In experiments 2 and 3, there was a suggestion that subjects’ tendency to switch their choices from one trial to the next was increased following anxiety induction and the recent occurrence of a panic attack, respectively. These findings were not hypothesised and effect sizes were small (and somewhat inconsistent across analysis methods), but given their consistency with a prior independent study (21), they warrant brief discussion. One possibility is that this increase in choice switching might reflect the enhanced uncertainty characteristic of anxious states (60) and could arise as a result of activation of the noradrenergic system (61, 62). Evidence for this comes from work suggesting that tonic noradrenaline release is linked to an increase in task irrelevant processing and a tendency to favour exploration over exploitation (63), characterised by some as a network ‘reset’ (64). This interpretation is limited by the absence of data on cortisol and noradrenaline response and the exploratory nature of the findings. Future research will be needed to test this more directly, using a cognitive test designed to explicitly separate exploration and exploitation.

This study had limitations. Firstly, null results are difficult to draw firm conclusions from. However, the findings of Experiment 3, which benefit from the inclusion of previously published clinical effect size comparator (the effect of compulsivity on model-based planning), help put these null findings in meaningful context. It is unlikely that our manipulation was not strong enough to induce a robust anxiogenic effect because previous studies have demonstrated that the 7.5% CO_2_ manipulation is powerful enough to elicit robust effects on behavioural performance relating to threat sensitivity and hyper-vigilance (33-35), in addition to the well-documented physiological and psychological effects (24, 58). The magnitude of self-report and physiological changes in the present study were on-par with those observed in prior studies (33-35). Finally, Bayesian analyses are presented in the Supplement and detail the extent to which evidence was in favour of the null, and this was in most cases in the ‘very strong’ range. A second limitation is that using panic attacks to measure ‘real world’ state anxiety is an imperfect methodology. Although panic attacks are associated with an increase in state anxiety (31), they are also associated with, and defined by, a much broader cascade of physical symptoms than the experience of state anxiety. However, this approach has two advantages over measuring self-reported state anxiety (e.g. using the STAI-state scale(65)). First, in the absence of an acute event (anxiety trigger), trait and state anxiety scores tend to be highly correlated (e.g. r=.71 (40)) and is thus thought to be more reflective of trait than state anxiety. Second, leveraging naturally occurring panic attacks allowed us to mirror the acute and sudden onset of anxiety that our lab-based procedure achieved.

## Conclusions

Experimentally-induced state anxiety failed to produce deficits in goal-directed behaviour as measured via two independent experiments using two well-validated probes. Such lack of effect was also observed in a more ecologically valid set-up, where we used recent panic attacks as a proxy for acute anxiety. While modest decreases in goal-directed planning were seen in individuals who had recent panic attacks in the past year, these effects did not survive when controlling for compulsivity. The same was true of the occurrence of major life stressors in the past year. From this we can conclude that it is unlikely that the deficits in goal-directed behaviour routinely identified in association with compulsivity are due to anxiety. More broadly, these data highlight the necessity of using positive clinical control measures and causal manipulations to ascertain robust and specific associations given a deeply complex and highly inter-correlated mental health landscape.

## Method

### Experiment 1

#### Subjects

88 participants were recruited through university mailing lists, departmental research panels and posted flyers within the University of Cambridge and the wider community. Experimental procedures were approved by the Ethics Committee of the University of Cambridge, School of the Biological Sciences. Exclusion criteria were screened by a structured telephone interview (Mini International Neuropsychiatric Interview: MINI(66)) and were as follows: current or past diagnosis of cardiovascular disease, respiratory disease, thyroid disease, or diabetes; lifetime history of DSM-VI Axis I disorders; having a first-degree relative diagnosed with panic disorder; (history of) migraine or epilepsy; pregnancy; excessive weekly consumption of alcohol (28 units for males, 21 units for females), excessive daily consumption of caffeine (more than 8 caffeinated drinks per day); current (illegal) drug use; recent history of smoking on a daily basis. Participants were free of regular medication intake, with the exception of the oral contraceptive pill. Invited participants were asked to abstain from alcohol consumption 24 hours prior to the experiment, as well as caffeinated drinks from the midnight before the experiment. Participants were randomly assigned to either the CO_2_-induced anxiety group (n= 43, 20 females; mean age = 27.55, SD = 11.04) or the normal air ‘placebo’ group (n= 45, 24 females; mean age = 27.40, SD = 10.03). Prior experimental work that was sensitive to identify a between-group effect of stress on habits employed samples of 51 (17) and 67 (18), respectively. We were considerably more conservative, testing 88 subjects.

#### Anxiety Manipulation

Participants were randomly assigned to two groups, one received the anxiety induction, which consisted of the inhalation of air enriched with 7.5 % CO_2_ (7.5% CO_2_, 20% O2, 71.5% N2, pre-mixed, BOC Special Gases, Guildford, UK) and one served as the control group, inhaling normal air. CO_2_ was administered in a single-blind manner while measuring goal-directed/habit behaviour via controlled tasks, and was designed to induce a physiological state of acute anxiety in a reliable and controlled manner (24). To measure the effectiveness of this procedure at inducing acute anxiety, we recorded physiological measurements comprising heart rate, diastolic and systolic blood pressure and psychological measurements comprising the 17-item Acute Panic Inventory (API: 67), 10-item Positive and Negative Affective Scale (PANAS(68)), and three Visual Analogue Scales assessing anxiety, fear, and happiness. Physiological measures were collected 10 minutes before, during and 15 minutes after the experimental manipulation. Psychological measures of subjective feeling due to the experimental manipulation were concomitantly collected, the only difference being that they were not interrogated during the performance of the task but immediately after and retrospectively on how they were feeling.

#### Contingency degradation manipulation

In a between-subjects design, participants were tested on the ability to detect action-outcome instrumental contingency via the experimental manipulation of contingency degradation. Our index of contingency was the standard ΔP measure indexing the action-outcome instrumental relationship (2). ΔP was the difference between the conditional probability of outcome given an action [P(O|A)], i.e., the probability of response-contingent outcome; and the probability of receiving an outcome given the absence of an action [P(O|-A)], i.e., the probability of a non-contingent outcome, such that ΔP = P(O|A) – P(O|-A). By increasing non-contingent outcomes, the contingency (i.e., the causal action-outcome association) is degraded. Under these circumstances, individuals who are making decisions in a goal-directed manner should stop or reduce responding in line with the reduction in instrumental contingency.

#### Contingency degradation paradigm

Subjects performed a contingency degradation task described previously (7) (Figure 1A). In a free-operant, self-paced procedure, a white triangle on the screen signalled that the participant was free to press, or not to press, the space bar. On each response, the triangle turned yellow until the end of the *a priori* specified bin to signal that a response has been recorded and to prevent multiple responses within the same 1-second bin. When a reward was delivered, following a key press or not, a 25 pence image was shown at the end of the bin for 500 milliseconds with the text “Reward, you win!” and a tone. If no outcome was delivered, no feedback was given and the next bin started. Each participant completed 8 blocks where ΔP was systematically varied (Figure 1B). A running total of money earned within the block was displayed in the corner of the screen and reset to 0 at the beginning of each block. Causality judgments regarding the relationship between pressing the key and receiving the reward were collected at the end of each block. Each block included 140 un-signalled bins, each lasting 1 second. If a response occurred during a given bin, the outcome was delivered with probability P(O|A) defined a priori for that block; if no response occurred, the outcome was delivered with probability P(O|-A) defined a priori for that block. Only the first space-bar press within the bin had any programmed consequences. By varying P(O|A) and P(O|-A), different levels of instrumental contingency were established in each block. In the first 2 blocks, all participants inhaled normal air and the associated programmed contingencies were always presented in the same order (high contingency 0.6, followed by degradation of the contingency to 0.3), providing an implicit training phase. The remaining blocks (test phase) were presented according to a Latin square design for participants in each of the two experimental groups.

#### Experienced contingency

As expected, for normal and CO_2_-enriched air condition, experienced contingencies (based on experienced event frequencies, see **Supplementary Material, Table S1**) matched the a priori programmed ones (CO_2_: r = 1.00, *p* < .001; Air: r = 1.00, *p* < .001). Therefore, programmed contingencies were used for subsequent analysis. Our findings were not confounded by between-group differences in experienced contingencies, as no main effect of group (*F*_(1, 63)_ = 0.80, *p* = .37) nor interaction between group and block (*F*_(2.57, 161.61)_ = 0.17, *p* = .89) was found.

#### Data analysis

We first performed analyses of variance (ANOVA) to determine whether there was a between-group difference in sensitivity to instrumental contingency as measured by response rate and causality judgment. Response rate was computed by dividing the number of bins for which a response was made by the total number of bins within each block. For each dependent variable, programmed contingency was used as a within-subject factor, and group was used as a between-subject factor. Analyses were conducted separately for the initial learning blocks and the test blocks. For the test blocks, we also investigated the relationship between response rate and contingency judgments, using a linear mixed-effects model. Specifically, we used contingency judgement and group as fixed effects, and we allowed the intercept and slope to vary between participants as random effects. We obtained p-values for the fixed effects using the Kenward-Roger method. Bayes factor analysis was used in case of failure to reject the null hypothesis, to examine the relative evidence for the null (results presented in the online supplement). We used the default JZS prior for the ANOVA models (69), and weakly informative priors for the mixed-effects model. Specifically, we used normal priors with mean=0 and standard deviation=10 for the fixed effect parameters; half student-t priors with degrees of freedom=3, location=0 and scale=10 for the standard deviation of the random effects; and an LKJ prior with shape=1 for the correlation between random effects. Analyses were performed in R version 3.4.3 (R Foundation for Statistical Computing, Vienna, Austria; http://www.r-project.org/) using the ‘afex’ package for ANOVA and linear mixed models, the ‘Bayes Factor’ and ‘brms’ package for Bayes factor analysis and the ‘tidyverse’ packages for data organization and visualization.

### Experiment 2

#### Subjects

61 healthy volunteers were recruited from the local community in the same manner as described in Experiment 1. Screening and exclusion criteria were identical to Experiment 1. Four subjects aborted the experiment and data were lost from an additional 3. Because of the nature of the analyses, subjects were excluded if their stay/switch behaviour showed such little variation as to preclude a hierarchical model-fit (choosing same response >90% of trials, N=3) or conversely, deviated substantially (>3 SDs) from the mean in the opposite direction (N=1). The final sample size for analysis was 50 (26 female) with ages ranging from 18-62. Sample size was determined based on previous study that detected a within-subjects (l-Dopa) effect on model-based planning using N=18 on this same task (38). As in Experiment 1, we were conservative in recruiting N=50. The study was approved by the same ethics committee as above.

#### Reinforcement learning task (37)

On each trial, participants were presented with a choice between two fractals, each of which commonly (70%; see Figure 3A) led to a particular second state displaying another two fractals. These second-state fractals each had some probability (between .25 and .75) of being rewarded with a pound coin. On 30 % of trials (“rare” transition trials; Figure 3A), choices uncharacteristically led to the other state. A purely model-free learner makes choices irrespective of these contingencies (i.e. which action is most strongly linked to which second stage state), and instead focuses on repeating actions that were followed by reward. A model-based strategy, in contrast, is characterized by sensitivity to both reward and the transition structure (contingency) within the task. This means that when a stage 1 action is ultimately rewarded at the end of a trial, a model-based learner will repeat that stage 1 action again, only if the path to reward they took was likely (i.e. involved a common transition). If the path they took to reward was unlikely (involving a rare transition), a model-based subject switches their stage 1 action to promote their chances of returning to that valuable second stage state. The chances that a second stage fractal would be rewarded drifted slowly over time, such that in order to perform optimally, subjects needed to update action preferences dynamically throughout the task.

Before starting the task, participants completed a training session, which comprised written instructions, the viewing of 20 trials demonstrating the probabilistic association between the second stage fractals and coin rewards, and completion of 20 trials of active practice with the probabilistic transition structure of the task. Subjects were then tested for their comprehension of the task with a short quiz (10) and if they answered any questions incorrectly, these comprehension issues were clarified on-screen. The task consisted of 200 trials in which participants had 2.5 s in which to make a response using the left and right keys following presentation of the first-state choice. If no response was made on time, “no response” were presented on the screen, and the next trial started. If a choice was made, the selected fractal moved to the top centre of the screen and shrunk in size. A new, second-state fractal appeared in the centre of the screen and was followed by a pound coin or a zero. Subjects completed two counterbalanced versions of the task, with different fractal stimuli and reward drifts.

#### Anxiety Induction

The anxiety induction procedure as well as collection of physiological and psychological measures was identical to Experiment 1, except for the within-subjects design. Participants attended a single test session during which they completed two versions of the Reinforcement Learning Task *during* 20min inhalation of air enriched with 7.5 % CO_2_ and normal air. Gas was administered in a single-blind manner and the order of CO_2_ versus air was counterbalanced.

#### Data Analysis

Data were analysed using mixed-effects logistic regression in the *lme4* package in R 3.5.1 (http://cran.us.r-project.org). In line with previous studies (3), we tested the extent to which subjects in general tend to repeat actions performed on the previous trial or explore a new one (‘Stay’: coded switch= 0; stay= 1), and whether these choices were influenced by whether or not their previous action was rewarded (‘Reward’: coded as rewarded = 1; unrewarded = −1), was followed by a rare or common transition (‘Transition’: coded as common= 1, rare= −1), and their interaction (‘Reward × Transition’). The intercept reflects tendencies to repeat the same action from one trial to the next, the main effect of reward reflects the contribution of model-free learning to subjects’ choices, while an interaction between Reward and Transition is the hallmark of model-based (goal-directed) behaviour. Ae included the anxiety induction as a within-subjects factor (coded CO_2_=1, Air=−1). We used Bound Optimization by Quadratic Approximation (bobyqa) with 1e5 functional evaluations. The model was specified as follows: Stay ∼ Reward*Transition*CO_2_ + (Reward* Transition*CO_2_ + 1|Subject). Bayes factor analysis was used in case of failure to reject the null hypothesis. We extracted estimates for model-based planning separately for each subject in each condition and used these to compare an ANOVA model with a within-subjects effect of gas to an intercept-only model. We also computed the model described above with self-reported sensitivity to the CO_2_ manipulation as an additional covariate, operationalized as change in self-reported Anxiety from Air to CO_2_.

#### Computational Modelling

A more elaborated form of this analysis is presented in the online supplement. In brief, this method allows for analysis of a greater number of potential behavioural confounds, including separating the distinct role of learning rate and choice randomness from that of model-based, model-free and choice repetition estimates from the simpler analysis. These results largely recapitulate the main findings of the paper, with slight differences flagged as appropriate.

### Experiment 3

#### Participants

Data were collected online using Amazon’s Mechanical Turk. Details of the experimental procedure can be found elsewhere, but in brief, data were analysed from 1,413 individuals (823 female) with ages ranging from 18 to 76 (M=33, SD=11), who were based in the USA, had a history of good performance (i.e. being paid in full on at least 95% of their previous tasks)). They were paid a base rate of $2.50, in addition to a bonus based on their earnings during the reinforcement-learning task (M=$0.54, SD=0.04). This study was approved by the New York University Committee on Activities Involving Human Subjects. These participants are the same as those in a previously published article (10).

#### Reinforcement learning task

The task employed in this study was the same as that described in Experiment 2. The only difference was that subjects completed it remotely, and that a more rigorous quality control procedure was implemented appropriate to online testing (detailed in Supplement).

#### Panic Attacks

The occurrence of recent panic attacks was assessed using item 1 on the self-report version of the Panic Disorder Severity Scale (PDSS, 41): “How many panic and limited symptoms attacks did you have during the week?”. Subjects were provided with a definition of a panic attack: a “sudden rush of fear or discomfort”, peaking within 10 minutes accompanied by 4 of 17 symptoms (e.g. rapid or pounding heartbeat, feeling of choking, nausea, chills or hot flushes, fear of dying). Subjects were told that episodes that have fewer than 4 symptoms are ‘limited symptom attacks’. Panic attack frequency scores ranged from none (“no panic or limited symptom attacks”), mild (no full panic attacks and no more than 1 limited symptom attack/day), moderate (“1 or 2 full panic attacks and/or multiple limited symptom attacks/day”), severe (Severe: more than 2 full attacks but not more than 1/day on average) and extreme (“full panic attacks occurred more than once a day, more days than not”).

#### Life Stress

Life stress was assessed using the Social Readjustment Scale (45), which presents an inventory of common stressful life events to participants and asks them to select those that applied to them in the previous 12 months. Events are weighted in a manner that reflects the relative amount of stress that event causes, with the death of a spouse and divorce being the most stressful and minor violations of the law, major holidays and vacations being the least. The present sample had a mean score of 159 (SD=120). Scores lower than 150 are considered evidence of ‘no significant stress’ (N=775), while scores in excess of 300 are considered signs of major stress (N=179 in this sample) (Figure 5B).

#### Control Variables

As detailed in a prior report, subjects completed a range of self-report questionnaires that were the topic of a factor analysis in a previously published study (10), which was subsequently validated in an independent dataset (70). One factor, titled “Compulsive Behaviour and Intrusive Thought”, was shown to be highly associated with model-based planning failures in this sample. Scores on this factor were thus controlled, along with IQ, age and gender.

#### Data Analysis

We performed the same analysis as in Experiment 2, but here we additionally controlled for variables that have been previously linked to model-based planning, namely: IQ, age, gender and a trans-diagnostic psychiatric trait “Compulsive Behaviour and Intrusive Thought”. Bayes factor analysis was conducted on a linear model where residuals for model-based planning was the dependent measure and life stress or panic symptoms were the experimental models compared to an intercept-only model. As in experiment 2, we complemented our regression analysis with a computational model, details of which are available in the online supplement.

## Acknowledgements

Research was funded by a Wellcome Trust Senior Investigator Award (TW Robbins 106431/Z/14/Z) and a Sir Henry Wellcome Postdoctoral Fellowship (CM Gillan 101521/Z/12/Z). CM Gillan is supported by a fellowship from MQ: transforming mental health (MQ16IP13). AB was supported by a fellowship from the Swiss National Science Foundation (SNF PASMP3-145749).

## References

1. Dickinson A. Actions and Habits: The Development of Behavioural Autonomy. Philosophical Transactions of the Royal Society of London Series B, Biological Sciences. 1985;308(1135):67–78.

2. Dickinson A, Balleine B. Motivational control of goal-directed action. Animal Learning & Behavior. 1994;22(1):1–18. doi: 10.3758/BF03199951. PubMed PMID: WOS:A1994NP64700001.

3. Daw ND, Gershman SJ, Seymour B, Dayan P, Dolan RJ. Model-Based Influences on Humans’ Choices and Striatal Prediction Errors. Neuron. 2011;69(6). doi: 10.1016/j.neuron.2011.02.027. PubMed PMID: WOS:000288886900015.

4. Gillan CM, Otto AR, Phelps EA, Daw ND. Model-based learning protects against forming habits. Cogn Affect Behav Neurosci. 2015. Epub 24/3/2015. doi: 10.3758/s13415-015-0347-6. PubMed PMID: 25801925.

5. Gillan CM, Papmeyer M, Morein-Zamir S, Sahakian BJ, Fineberg NA, Robbins TW, et al. Disruption in the balance between goal-directed behavior and habit learning in obsessive-compulsive disorder. Am J Psychiatry. 2011;168(7):718–26. Epub 2011/05/17. doi: appi.ajp.2011.10071062 [pii] 10.1176/appi.ajp.2011.10071062. PubMed PMID: 21572165.

6. Gillan CM, Robbins TW. Goal-directed learning and obsessive-compulsive disorder. Philos Trans R Soc Lond B Biol Sci. 2014;369(1655). doi: 10.1098/rstb.2013.0475. PubMed PMID: 25267818; PubMed Central PMCID: PMCPMC4186229.

7. Vaghi MM, Cardinal RN, Apergis-Schoute AM, Fineberg NA, Sule A, Robbins TW. Action-Outcome Knowledge Dissociates From Behavior in Obsessive-Compulsive Disorder Following Contingency Degradation. Biol Psychiatry Cogn Neurosci Neuroimaging. 2018. Epub 2018/10/09. doi: 10.1016/j.bpsc.2018.09.014. PubMed PMID: 30545754.

8. Voon V, Derbyshire K, Rück C, Irvine MA, Worbe Y, Enander J, et al. Disorders of compulsivity: a common bias towards learning habits. Mol Psychiatry. 2014. doi: 10.1038/mp.2014.44. PubMed PMID: 24840709.

9. Sjoerds Z, de Wit S, van den Brink W, Robbins TW, Beekman AT, Penninx BW, et al. Behavioral and neuroimaging evidence for overreliance on habit learning in alcohol-dependent patients. Transl Psychiatry. 2013;3:e337. doi: 10.1038/tp.2013.107. PubMed PMID: 24346135; PubMed Central PMCID: PMCPMC4030326.

10. Gillan C, Kosinski M, Whelan R, Phelps E, Daw N. Characterizing a psychiatric symptom dimension related to deficits in goal-directed control. eLife. 2016;5(e11305). doi: http://dx.doi.org/10.7554/eLife.11305.

11. Robbins TW, Gillan CM, Smith DG, de Wit S, Ersche KD. Neurocognitive endophenotypes of impulsivity and compulsivity: towards dimensional psychiatry. Trends Cogn Sci. 2012;16(1):81–91. doi: 10.1016/j.tics.2011.11.009. PubMed PMID: 22155014.

12. Nestadt G, Di CZ, Riddle MA, Grados MA, Greenberg BD, Fyer AJ, et al. Obsessive-compulsive disorder: subclassification based on co-morbidity. Psychol Med. 2009;39(9):1491–501. doi: 10.1017/S0033291708004753. PubMed PMID: 19046474; PubMed Central PMCID: PMCPMC3039126.

13. Stein DJ, Fineberg NA, Bienvenu OJ, Denys D, Lochner C, Nestadt G, et al. Should OCD be classified as an anxiety disorder in DSM-V? Depress Anxiety. 2010;27(6):495–506. Epub 2010/06/10. doi: 10.1002/da.20699. PubMed PMID: 20533366.

14. Shields GS, Sazma MA, Yonelinas AP. The effects of acute stress on core executive functions: A meta-analysis and comparison with cortisol. Neurosci Biobehav Rev. 2016;68:651–68. Epub 2016/06/28. doi: 10.1016/j.neubiorev.2016.06.038. PubMed PMID: 27371161; PubMed Central PMCID: PMCPMC5003767.

15. Alvares GA, Balleine BW, Guastella AJ. Impairments in goal-directed actions predict treatment response to cognitive-behavioral therapy in social anxiety disorder. PLoS One. 2014;9(4):e94778. doi: 10.1371/journal.pone.0094778. PubMed PMID: 24728288; PubMed Central PMCID: PMCPMC3984205.

16. Mataix-Cols D, Cullen S, Lange K, Zelaya F, Andrew C, Amaro E, et al. Neural correlates of anxiety associated with obsessive-compulsive symptom dimensions in normal volunteers. Biological Psychiatry. 2003;53(6):482–93. doi: 10.1016/s0006-3223(03)01504-4. PubMed PMID: WOS:000181569900003.

17. Schwabe L, Wolf OT. Stress prompts habit behavior in humans. J Neurosci. 2009;29(22):7191–8. Epub 2009/06/06. doi: 29/22/7191 [pii] 10.1523/JNEUROSCI.0979-09.2009. PubMed PMID: 19494141.

18. Schwabe L, Wolf OT. Socially evaluated cold pressor stress after instrumental learning favors habits over goal-directed action. Psychoneuroendocrinology. 2010. Epub 2010/01/15. doi: S0306-4530(09)00372-2 [pii] 10.1016/j.psyneuen.2009.12.010. PubMed PMID: 20071096.

19. Park H, Lee D, Chey J. Stress enhances model-free reinforcement learning only after negative outcome. PLoS One. 2017;12(7):e0180588. Epub 2017/07/19. doi: 10.1371/journal.pone.0180588. PubMed PMID: 28723943; PubMed Central PMCID: PMCPMC5516979.

20. Otto AR, Raio CM, Chiang A, Phelps EA, Daw ND. Working-memory capacity protects model-based learning from stress. Proc Natl Acad Sci U S A. 2013;110(52):20941–6. doi: 10.1073/pnas.1312011110. PubMed PMID: 24324166; PubMed Central PMCID: PMCPMC3876216.

21. Radenbach C, Reiter AM, Engert V, Sjoerds Z, Villringer A, Heinze HJ, et al. The interaction of acute and chronic stress impairs model-based behavioral control. Psychoneuroendocrinology. 2015;53:268–80. Epub 2015/01/07. doi: 10.1016/j.psyneuen.2014.12.017. PubMed PMID: 25662093.

22. Heller AS, Ezie CEC, Otto AR, Timpano KR. Model-based learning and individual differences in depression: The moderating role of stress. Behav Res Ther. 2018;111:19–26. Epub 2018/09/26. doi: 10.1016/j.brat.2018.09.007. PubMed PMID: 30273768.

23. Shin LM, Liberzon I. The neurocircuitry of fear, stress, and anxiety disorders. Neuropsychopharmacology. 2010;35(1):169–91. doi: 10.1038/npp.2009.83. PubMed PMID: 19625997; PubMed Central PMCID: PMCPMC3055419.

24. Bailey JE, Argyropoulos SV, Kendrick AH, Nutt DJ. Behavioral and cardiovascular effects of 7.5% CO2 in human volunteers. Depress Anxiety. 2005;21(1):18–25. doi: 10.1002/da.20048. PubMed PMID: 15782425.

25. Woods SW, Charney DS, Goodman WK, Heninger GR. Carbon dioxide-induced anxiety. Behavioral, physiologic, and biochemical effects of carbon dioxide in patients with panic disorders and healthy subjects. Arch Gen Psychiatry. 1988;45(1):43–52. PubMed PMID: 3122696.

26. Perna G, Barbini B, Cocchi S, Bertani A, Gasperini M. 35% CO2 challenge in panic and mood disorders. J Affect Disord. 1995;33(3):189–94. PubMed PMID: 7790671.

27. Griez E, Zandbergen J, Pols H, de Loof C. Response to 35% CO2 as a marker of panic in severe anxiety. Am J Psychiatry. 1990;147(6):796–7. doi: 10.1176/ajp.147.6.796. PubMed PMID: 2111639.

28. Argyropoulos SV, Bailey JE, Hood SD, Kendrick AH, Rich AS, Laszlo G, et al. Inhalation of 35% CO(2) results in activation of the HPA axis in healthy volunteers. Psychoneuroendocrinology. 2002;27(6):715–29. PubMed PMID: 12084664.

29. Perna G, Bertani A, Caldirola D, Bellodi L. Family history of panic disorder and hypersensitivity to CO2 in patients with panic disorder. Am J Psychiatry. 1996;153(8):1060–4. doi: 10.1176/ajp.153.8.1060. PubMed PMID: 8678175.

30. Perna G, Battaglia M, Garberi A, Arancio C, Bertani A, Bellodi L. Carbon dioxide/oxygen challenge test in panic disorder. Psychiatry Res. 1994;52(2):159–71. PubMed PMID: 7972572.

31. Aronson TA, Logue CM. Phenomenology of panic attacks: a descriptive study of panic disorder patients’ self-reports. J Clin Psychiatry. 1988;49(1):8–13. PubMed PMID: 3335492.

32. Vyas A, Pillai AG, Chattarji S. Recovery after chronic stress fails to reverse amygdaloid neuronal hypertrophy and enhanced anxiety-like behavior. Neuroscience. 2004;128(4):667–73. doi: 10.1016/j.neuroscience.2004.07.013. PubMed PMID: 15464275.

33. Garner M, Attwood A, Baldwin DS, Munafò MR. Inhalation of 7.5% carbon dioxide increases alerting and orienting attention network function. Psychopharmacology (Berl). 2012;223(1):67–73. Epub 2012/03/29. doi: 10.1007/s00213-012-2690-4. PubMed PMID: 22453547.

34. Cooper R, Howard CJ, Attwood AS, Stirland R, Rostant V, Renton L, et al. Acutely induced anxiety increases negative interpretations of events in a closed-circuit television monitoring task. Cogn Emot. 2013;27(2):273–82. Epub 2012/07/11. doi: 10.1080/02699931.2012.704352. PubMed PMID: 22780582.

35. Garner M, Attwood A, Baldwin DS, James A, Munafò MR. Inhalation of 7.5% carbon dioxide increases threat processing in humans. Neuropsychopharmacology. 2011;36(8):1557–62. Epub 2011/04/13. doi: 10.1038/npp.2011.15. PubMed PMID: 21490591; PubMed Central PMCID: PMCPMC3138667.

36. Dickinson A, Nicholas DJ, Adams CD. The effect of the instrumental training contingency on susceptibility to reinforcer devaluation. Quarterly Journal of Experimental Psychology Section B-Comparative and Physiological Psychology. 1983;35(FEB). PubMed PMID: WOS:A1983QE08400003.

37. Daw ND, Niv Y, Dayan P. Uncertainty-based competition between prefrontal and dorsolateral striatal systems for behavioral control. Nature Neuroscience. 2005;8(12). doi: 10.1038/nn1560. PubMed PMID: WOS:000233576200019.

38. Wunderlich K, Smittenaar P, Dolan RJ. Dopamine Enhances Model-Based over Model-Free Choice Behavior. Neuron. 2012;75(3). doi: 10.1016/j.neuron.2012.03.042. PubMed PMID: WOS:000307417700010.

39. Worbe Y, Palminteri S, Savulich G, Daw ND, Fernandez-Egea E, Robbins TW, et al. Valence-dependent influence of serotonin depletion on model-based choice strategy. Mol Psychiatry. 2015. doi: 10.1038/mp.2015.46. PubMed PMID: 25869808; PubMed Central PMCID: PMCPMC4519524.

40. Grös DF, Antony MM, Simms LJ, McCabe RE. Psychometric properties of the State-Trait Inventory for Cognitive and Somatic Anxiety (STICSA): comparison to the State-Trait Anxiety Inventory (STAI). Psychol Assess. 2007;19(4):369–81. doi: 10.1037/1040-3590.19.4.369. PubMed PMID: 18085930.

41. Shear MK, Brown TA, Barlow DH, Money R, Sholomskas DE, Woods SW, et al. Multicenter collaborative panic disorder severity scale. Am J Psychiatry. 1997;154(11):1571–5. doi: 10.1176/ajp.154.11.1571. PubMed PMID: 9356566.

42. Barrera TL, Wilson KP, Norton PJ. The experience of panic symptoms across racial groups in a student sample. J Anxiety Disord. 2010;24(8):873–8. Epub 2010/06/19. doi: 10.1016/j.janxdis.2010.06.010. PubMed PMID: 20621442; PubMed Central PMCID: PMCPMC2956784.

43. Reed V, Wittchen HU. DSM-IV panic attacks and panic disorder in a community sample of adolescents and young adults: how specific are panic attacks? J Psychiatr Res. 1998;32(6):335–45. PubMed PMID: 9844949.

44. Goodwin RD, Lieb R, Hoefler M, Pfister H, Bittner A, Beesdo K, et al. Panic attack as a risk factor for severe psychopathology. Am J Psychiatry. 2004;161(12):2207–14. doi: 10.1176/appi.ajp.161.12.2207. PubMed PMID: 15569891.

45. Holmes TH, Rahe RH. The Social Readjustment Rating Scale. J Psychosom Res. 1967;11(2):213–8. PubMed PMID: 6059863.

46. Mogg K, Bradley BP, de Bono J, Painter M. Time course of attentional bias for threat information in non-clinical anxiety. Behav Res Ther. 1997;35(4):297–303. PubMed PMID: 9134784.

47. Eysenck MW, Mogg K, May J, Richards A, Mathews A. Bias in interpretation of ambiguous sentences related to threat in anxiety. J Abnorm Psychol. 1991;100(2):144–50. PubMed PMID: 2040764.

48. Paulus MP, Yu AJ. Emotion and decision-making: affect-driven belief systems in anxiety and depression. Trends Cogn Sci. 2012;16(9):476–83. Epub 2012/08/13. doi: 10.1016/j.tics.2012.07.009. PubMed PMID: 22898207; PubMed Central PMCID: PMCPMC3446252.

49. Mueller EM, Nguyen J, Ray WJ, Borkovec TD. Future-oriented decision-making in Generalized Anxiety Disorder is evident across different versions of the Iowa Gambling Task. J Behav Ther Exp Psychiatry. 2010;41(2):165–71. Epub 2009/12/28. doi: 10.1016/j.jbtep.2009.12.002. PubMed PMID: 20060098.

50. Miu AC, Heilman RM, Houser D. Anxiety impairs decision-making: psychophysiological evidence from an Iowa Gambling Task. Biol Psychol. 2008;77(3):353–8. Epub 2007/12/07. doi: 10.1016/j.biopsycho.2007.11.010. PubMed PMID: 18191013.

51. Dias-Ferreira E, Sousa JC, Melo I, Morgado P, Mesquita AR, Cerqueira JJ, et al. Chronic stress causes frontostriatal reorganization and affects decision-making. Science. 2009;325(5940):621–5. Epub 2009/08/01. doi: 325/5940/621 [pii] 10.1126/science.1171203. PubMed PMID: 19644122.

52. Schwabe L, Haddad L, Schachinger H. HPA axis activation by a socially evaluated cold-pressor test. Psychoneuroendocrinology. 2008;33(6):890–5. Epub 2008/04/09. doi: 10.1016/j.psyneuen.2008.03.001. PubMed PMID: 18403130.

53. Schwabe L, Tegenthoff M, Höffken O, Wolf OT. Concurrent glucocorticoid and noradrenergic activity shifts instrumental behavior from goal-directed to habitual control. J Neurosci. 2010;30(24):8190–6. doi: 10.1523/JNEUROSCI.0734-10.2010. PubMed PMID: 20554869.

54. Schwabe L, Tegenthoff M, Höffken O, Wolf OT. Simultaneous glucocorticoid and noradrenergic activity disrupts the neural basis of goal-directed action in the human brain. J Neurosci. 2012;32(30):10146–55. doi: 10.1523/JNEUROSCI.1304-12.2012. PubMed PMID: 22836250.

55. Bailey JE, Argyropoulos SV, Lightman SL, Nutt DJ. Does the brain noradrenaline network mediate the effects of the CO2 challenge? J Psychopharmacol. 2003;17(3):252–9. doi: 10.1177/02698811030173002. PubMed PMID: 14513913.

56. Allen AP, Kennedy PJ, Cryan JF, Dinan TG, Clarke G. Biological and psychological markers of stress in humans: focus on the Trier Social Stress Test. Neurosci Biobehav Rev. 2014;38:94–124. Epub 2013/11/14. doi: 10.1016/j.neubiorev.2013.11.005. PubMed PMID: 24239854.

57. Oliveira D, Chagas M, Garcia L, Crippa J, Zuardi A. Oxytocin interference in the effects induced by inhalation of 7.5% CO 2 in healthy volunteers. Hum Psychopharmacol Clin Exp. 2012;27 378–85.

58. Bailey JE, Kendrick A, Diaper A, Potokar JP, Nutt DJ. A validation of the 7.5% CO2 model of GAD using paroxetine and lorazepam in healthy volunteers. J Psychopharmacol. 2007;21(1):42–9. Epub 2006/03/13. doi: 10.1177/0269881106063889. PubMed PMID: 16533865.

59. Diaper A, Papadopoulos A, Rich AS, Dawson GR, Dourish CT, Nutt DJ, et al. The effect of a clinically effective and non-effective dose of lorazepam on 7.5% CO2-induced anxiety. Hum Psychopharmacol. 2012;27(6):540–8. Epub 2012/10/02. doi: 10.1002/hup.2261. PubMed PMID: 23027657.

60. Grupe DW, Nitschke JB. Uncertainty and anticipation in anxiety: an integrated neurobiological and psychological perspective. Nat Rev Neurosci. 2013;14(7):488–501. doi: 10.1038/nrn3524. PubMed PMID: 23783199; PubMed Central PMCID: PMCPMC4276319.

61. Yu A, Dayan P. Uncertainty, neuromodulation, and attention. Neuron. 2005;46(4).

62. Redmond DE, Huang YH. Current concepts. II. New evidence for a locus coeruleus-norepinephrine connection with anxiety. Life Sci. 1979;25(26):2149–62. PubMed PMID: 120478.

63. Aston-Jones G, Cohen JD. An integrative theory of locus coeruleus-norepinephrine function: adaptive gain and optimal performance. Annu Rev Neurosci. 2005;28:403–50. doi: 10.1146/annurev.neuro.28.061604.135709. PubMed PMID: 16022602.

64. Bouret S, Sara SJ. Network reset: a simplified overarching theory of locus coeruleus noradrenaline function. Trends Neurosci. 2005;28(11):574–82. Epub 2005/09/13. doi: 10.1016/j.tins.2005.09.002. PubMed PMID: 16165227.

65. Spielberger CD, Gorsuch RL, Lushene R, Vagg PR, Jacobs GA. Manual for the State-Trait Anxiety Inventory. Palo Alto, CA: Consulting Psychologists Press; 1983.

66. Sheehan DV, Lecrubier Y, Sheehan KH, Amorim P, Janavs J, Weiller E, et al. The Mini-International Neuropsychiatric Interview (M.I.N.I.): the development and validation of a structured diagnostic psychiatric interview for DSM-IV and ICD-10. J Clin Psychiatry. 1998;59 (Suppl 20):22–33. PubMed PMID: 9881538.

67. Liebowitz MR, Fyer AJ, Gorman JM, et al. Lactate provocation of panic attacks: I. clinical and behavioral findings. Arch Gen Psychiatry. 1984;41(8):764–70.

68. Watson D, Clark LA, Tellegen A. Development and validation of brief measures of positive and negative affect: the PANAS scales. J Pers Soc Psychol. 1988;54(6):1063–70. PubMed PMID: 3397865.

69. Rouder JN, Speckman PL, Sun D, Morey RD, Iverson G. Bayesian t tests for accepting and rejecting the null hypothesis. Psychon Bull Rev. 2009;16(2):225–37. doi: 10.3758/PBR.16.2.225. PubMed PMID: 19293088.

70. Rouault M, Seow T, Gillan CM, Fleming SM. Psychiatric Symptom Dimensions Are Associated With Dissociable Shifts in Metacognition but Not Task Performance. Biol Psychiatry. 2018;84(6):443–51. Epub 2018/01/11. doi: 10.1016/j.biopsych.2017.12.017. PubMed PMID: 29458997; PubMed Central PMCID: PMCPMC6117452.

