## Supplementary Materials for "Experimentally-induced and real-world acute anxiety have no effect on goal-directed behaviour"

*Denotes equal first author contribution

**Experiment 1**

**Table S1: Programmed contingency, experienced contingency, response rate, and causality judgement for each experimental block.**

|  |  |  | **Programmed**  **contingency** | | | **Experienced contingency** | | **Response**  **rate** | | **Causality judgment** | |
| --- | --- | --- | --- | --- | --- | --- | --- | --- | --- | --- | --- |
|  | **Block** | | **P(O\|A)** | **P(O\|-A)** | **ΔP** | **Air** | **CO_2_** | **Air** | **CO_2_** | **Air** | **CO_2_** |
| **Fixed Order** | 1 | | 0.60 | 0.00 | 0.60 | 0.61 | 0.60 | 0.54 | 0.57 | 58.79 | 63.19 |
|  | 2 | | 0.60 | 0.30 | 0.30 | 0.36 | 0.35 | 0.42 | 0.46 | 40.15 | 43.84 |
| **Shuffled in a Latin square** | 3 | | 0.60 | 0.00 | 0.60 | 0.60 | 0.59 | 0.64 | 0.59 | 56.29 | 58.97 |
|  | 4 | | 0.60 | 0.10 | 0.50 | 0.52 | 0.51 | 0.55 | 0.55 | 47.53 | 51.61 |
|  | 5 | | 0.60 | 0.30 | 0.30 | 0.30 | 0.31 | 0.50 | 0.42 | 42.68 | 40.53 |
|  | 6 | | 0.60 | 0.40 | 0.20 | 0.21 | 0.19 | 0.40 | 0.45 | 38.65 | 40.22 |
|  | 7 | | 0.60 | 0.50 | 0.10 | 0.07 | 0.06 | 0.43 | 0.40 | 33.66 | 36.87 |
|  | 8 | | 0.60 | 0.60 | 0.0 | -0.02 | -0.03 | 0.39 | 0.33 | 28.27 | 28.10 |

**P(O|A), probability of the outcome given the action; P(O|-A), probability of the outcome in the absence of the action; ΔP =contingency. Dependent variables are given as mean. Blocks 1-2 were presented in a fixed order; Block 4-8 were presented according to a Latin square design. Programmed contingency refers to the a priori experimentally programmed contingency resulting from the a priori programmed conditional probabilities. Experienced contingency was computed on the basis of experienced event frequencies.**

**Psychological and physiological response to stress.** Psychological and physiological measures confirmed that participants in the CO_2_ condition experienced greater anxiety and stress than those assigned to the Air condition (Figure 1B and 1D). Group means and standard deviations are presented in the supplementary materials (Table S2 and S3, respectively). Subjective negative affect increased under CO_2_; there were significant group by time interaction effects (all *p* < .001) for the API, the PANAS negative affect subscale, as well as the “fearful” and “anxious” visual analogue scales (Figure 1D). The results for positive affect were mixed; happiness decreased under CO_2_ (*p* = .004), but there no significant difference for the (more extensive) PANAS positive affect subscale (*p* = .84). In terms of autonomic measures of arousal, there were significant group by time interaction effects (all *p* < .001) for heart rate, systolic blood pressure and diastolic blood pressure. As shown in Figure 1B, heart rate and blood pressure significantly increased under inhalation of CO_2_ compared to normal air.

**Relationship between response rate and causality judgments.** As goal-directed control involves the implementation of contingency knowledge into flexible action, we lastly tested the extent to which causality judgments predicted response rate, and whether that might be affected by CO_2_-induced anxiety. Overall, response rate was linearly predicted by causality ratings (*F*_(1, 50.51)_ = 78.84, *p* < .001), but the slope of this relationship was not significantly different between groups (group by causality judgement interaction effect: *F*_(1, 50.51)_ = 0.05, *p* = .83) (Figure 2B). This analysis thus indicated that the linear relationship between subjectively detected instrumental contingency and response rate remained intact in face of an acute anxiety induction.

**Bayes Factor analysis.** For the three key analysis models described above, Bayes factor analysis indicated that the null model was strongly preferred over the alternative model that included the acute anxiety manipulation. Specifically, for the analyses of the effect of instrumental contingency (ΔP) on response rate and causality judgement, the null model (including only contingency) was strongly preferred over the alternative model with a main effect of anxiety and interaction effect of anxiety by contingency (BF_01_ = 16.81 and BF_01_ = 386.15, respectively). Similarly, for the analysis of the relationship between response rate and causality judgments, the null model (including only causality judgements) was very strongly preferred over the alternative model with fixed effect of anxiety and anxiety by causality judgement interaction (BF_01_ = 5882.35).

**Table S2: Means and standard deviations for positive and negative affect by group and time**

|  | **Pretest** | | **Test** | | **Post-test** | |
| --- | --- | --- | --- | --- | --- | --- |
|  | **Air** | **CO_2_** | **Air** | **CO_2_** | **Air** | **CO_2_** |
| **API** | 1.80 ± 2.00 | 2.88 ± 3.28 | 2.98 ± 3.69 | 16.5 ± 8.17 | 3.10 ± 3.99 | 4.35 ± 6.05 |
| **VAS** |  |  |  |  |  |  |
| Anxious | 17.49± 16.95 | 18.19± 18.10 | 11.39± 11.77 | 42.34± 27.08 | 13.22± 14.05 | 16.83± 19.82 |
| Fearful | 14.48± 19.15 | 12.88± 16.22 | 10.23± 15.12 | 35.59± 27.52 | 11.84± 17.76 | 14.16± 19.70 |
| Happy | 63.23± 18.20 | 53.66± 19.96 | 54.26± 19.98 | 34.11± 19.62 | 59.05± 21.93 | 52.70± 22.69 |
| **PANAS** |  |  |  |  |  |  |
| Negative | 12.67 ± 3.27 | 12.71 ± 3.23 | 11.41 ± 2.24 | 17.28 ± 6.42 | 12.19 ± 3.48 | 12.65 ± 4.62 |
| Positive | 27.60 ± 7.76 | 26.95 ± 8.35 | 22.14 ± 8.30 | 20.67 ± 7.92 | 24.43 ± 9.29 | 23.47 ± 8.64 |

**API, Acute Panic Inventory; VAS, Visual Analogue Scale; PANAS, Positive and Negative Affective Scale. Data show mean and standard deviation.**

**Table S3: Means and standard deviations for autonomic arousal by group and time**

|  | **Pretest** | | **Test** | | **Post-test** | |
| --- | --- | --- | --- | --- | --- | --- |
|  | **Air** | **CO_2_** | **Air** | **CO_2_** | **Air** | **CO_2_** |
| **HR** | 70.0± 10.4 | 70.1± 10.5 | 70.3± 10.0 | 86.0± 15.8 | 68.1± 11.1 | 68.0± 13.2 |
| **BP systolic** | 116.0±13.6 | 117.7±16.3 | 115.0±12.3 | 140.4±22.8 | 115.2±20.1 | 123.3±16.7 |
| **BP diastolic** | 69.2± 8.2 | 71.6± 10.4 | 70.5 ± 9.9 | 83.7± 15.8 | 72.2 ± 9.7 | 78.5± 11.7 |

**HR, rate rate; BP, blood pressure. Data show mean and standard deviation.**

**Supplementary Information for Experiment 2**

**Psychological and physiological response to stress.** Under acute CO_2_ administration, subjects were more anxious, fearful, and less happy (all *p*<.001, Table 2). Subjects scores on the acute panic index also increased (*p*<.001) under CO_2_, as did their heart-rate (*p*=.002) and blood pressure (*p*<.001). Subjects also reported more negative affect (*p*<.001) and less positive affect (*p*=.029) on the PANAS.

**Table S4. Self-report within-subject changes associated with acute CO_2_ administration**

|  | **Air** | **CO_2_** | **F** | ***p*** |
| --- | --- | --- | --- | --- |
|  | *Mean (SD)* | *Mean (SD)* |  |  |
| **API** | 3.6 (4.0) | 13.4 (9.0) | 64.86 | < .001*** |
| **PANAS PA** | 24.6 (8.4) | 22.4 (8.9) | 5.16 | = .03* |
| **PANAS NA** | 11.5 (2.7) | 15.9 (5.1) | 55.1 | < .001*** |
| **VAS anxious** | 13.6 (12.6) | 35.3 (24.0) | 57.47 | < .001*** |
| **VAS fearful** | 9.4 (9.8) | 25.6 (24.1) | 35.44 | < .001*** |
| **VAS happy** | 51.5 (20.7) | 41.6 (22.2) | 15.72 | < .001*** |
| **BP-systolic^** | 114.8 (15.2) | 133.9 (22.2) | 74.17 | < .001*** |
| **BP-diastolic^** | 73.1(12.0) | 81.1(15.3) | 15.7 | < .001*** |
| **HR^** | 67.2 (8.9) | 75.1 (17.9) | 10.71 | =.002** |

**SD= standard deviation; API = acute panic index; PANAS= positive and negative affect schedule; PA= positive affect; NA= negative affect’ VAS= visual analogue scale; BP= blood pressure; HR= heart rate.**

**^Physiological data were missing from 1 subject (leaving N=49 for BP and HR measures).**

**** p<.05; **p<.01, ***p<.001***

**Detailed Results for Model-Based Task.** The regression model fit subjects’ behavior as expected, based on the prior literature; there was a significant main effect of Reward (β =.55, SE=.08, *p*<.001) and a significant Reward x Transition interaction (β=.28, SE=.06, *p*<.001), providing evidence that, overall, subjects’ choices showed signatures of both model-free and model-based processes. The intercept was significant; subjects had an overall tendency to repeat choices from one trial to the next, β=1.59, SE=.12, *p*<.001) (Table S5). Importantly, CO_2_ had no effect on subjects’ tendency to exhibit model-based (β=-0.03, SE=0.04, *p*=.44) or model-free (β=-0.02, SE=0.03, *p*=.52) behavior. There was a non-significant trend for subjects to switch more under CO_2_ (main effect of CO_2_ condition, β=-0.08, SE=0.04, *p*=.060; Table S5). Bayes factor analysis indicated that there was moderate evidence in favor of the null model over the alternative model that included the acute anxiety manipulation (BF_01_ = 3.5).

**Table S5. Results from regression model for Experiment 1**

| **Coefficient** | **β (SE)** | ***z*-value** | ***p*-value** |
| --- | --- | --- | --- |
| (Intercept) | 1.59(0.12) | 12.88 | <.001 *** |
| Reward | 0.55(0.08) | 6.74 | <.001 *** |
| Transition | 0.08(0.04) | 1.96 | 0.05 * |
| CO_2_ | -0.08(0.04) | -1.85 | 0.06 |
| Reward:Transition | 0.28(0.06) | 4.48 | <.001 *** |
| Reward:CO_2_ | -0.02(0.03) | -0.65 | 0.52 |
| Transition:CO_2_ | 0.04(0.03) | 1.48 | 0.14 |
| Reward:Transition:CO_2_ | -0.03(0.04) | -0.76 | 0.44 |

****p*<.05	** *p*<.01	****p*<.001**

**SE=standard error**

**Computational Modeling Method**

*Reinforcement Learning (RL) Model*

We used a reinforcement-learning (RL) model based on a hybrid of model-free Q_MF_(s_A_, a) and model-based Q_MB_(s_A_, a), as utilized in previous studies (1, 2). This model consists of separate model-based and model-free subcomponents, both of which estimate a state-action value function, which maps each possible action to its expected future reward. On trial t, we denote the first-stage state (always s_A_) by s_1,t_, the second-stage states by s_2,t_, the chosen first-stage action by a_t_, and the second-stage rewards as r_t_ .

For the model-free algorithm, we used temporal difference (TD) learning (3), which updates the value for the visited state-action pair at s_1,t_ according to: $Q_{MF}\left( s_{1,t},a_{t} \right)=Q_{MF}\left( s_{1,t},a_{t} \right)+\alpha\delta_{1,t}$

where α is a learning rate parameter and$\delta_{1,t}$ is the reward prediction error (RPE) at state 1, trial t:$\delta_{1,t}=Q_{MF}\left( s_{2,t} \right) {- Q}_{MF}\left( s_{1,t},a_{t} \right)$

The RPE is based on the second-stage value, $Q_{MF}\left( S_{2,t} \right).$Second-stage values are themselves updated according to: $Q_{MF}\left( s_{2,t} \right)=Q_{MF}\left( s_{2,t} \right)+\alpha\delta_{2,t}$

where the RPE at the second stage state, trial t $(\delta_{2,t})$ is determined by whether or not the trial was rewarded, $r_{t}$:$\delta_{2,t}=r_{t} {- Q}_{MF}\left( s_{2,t} \right)$

The model assumes that the eligibility trace =1 for all subjects (1), thus propagating second-stage reward information to the first-stage values. At the end of each trial, we decayed the Q values for all of the non-selected actions by multiplying them by 1 − α (4, 5).

The model-based RL algorithm works by learning the transition structure of the task (the state most often visited previously after each top-stage choice) and immediate reward values for each second stage state, then computing cumulative state-action values by iterative expectation over these. At the second stage (where immediate rewards were offered), the problem of learning immediate rewards is equivalent to that for TD above, because $Q_{MF}\left( s_{2t} \right)$ is just an estimate of the immediate reward r_t_; with no further stages to anticipate, and the SARSA learning rule reduces to a delta rule for predicting the immediate reward. Thus, the two approaches coincide at the second stage, and we define Q_MB_ = Q_MF_ at those states. Critically, the top level model-based values are defined from both the transition and reward estimates using the Bellman Equation (6):

$$Q_{MB}\left( s_{A}{,a}_{A_{j}} \right)={P\left( s_{B}|s_{A}{,a}_{j} \right) Q}_{MF}\left( s_{B} \right)+{P\left( s_{C}|s_{A}{,a}_{j} \right) Q}_{MF}\left( s_{C} \right)$$

where we have assumed these are recomputed on each trial from the current estimates of the transition probabilities and rewards. To connect the model-based and model-free values to choices, we use a softmax choice rule, which assigns a probability to each action based on a weighted sum of model-based and model free values (7). The probability of each choice at the first stage is calculated, accordingly, as

$$P\left( a_{t}= a|s_{1,t} \right)=\frac{{exp[\beta_{MB}\cdot Q}_{MB}\left( s_{1,t},a \right)+\beta_{MF}\cdot Q_{MF}\left( s_{1,t},a \right)+p\cdot rep(a)]}{\Sigma_{a^{'}} {exp[\beta_{MB}\cdot Q}_{MB}\left( s_{1,t},a' \right)+\beta_{MF}\cdot Q_{MF}\left( s_{1,t},a' \right)+p\cdot rep(a')]}$$

The indicator function rep($a$) is defined as 1 if $a$ is the same one as was chosen on the previous trial, zero otherwise. Together with the “stickiness” parameter p, this captures first-order perseveration (p > 0) or switching (p < 0) in the first- stage choices (4). Second-stage choices are modeled with only a single value term$Q_{MF}\left( s_{1,t},a \right)$ with its an inverse temperature β and no stickiness parameter.

This model was embedded within a multi-level random effects model of the population variation in its parameters to estimate it for all subjects simultaneously and to estimate the effect of condition on these parameters, i.e. CO_2_ (on/off). This was done identically to Sharp and colleagues (1), in that the within-subjects effect of CO_2_ is a subject-specific latent variable with its own population-level mean and variance, which are themselves inferred. All of the parameters of the model were taken as random effects, instantiated separately for each subject *s* from a common group level distribution. We estimated the parameters of the group level distributions using uninformative priors: for all parameters, the prior means and SDs were the heavy-tailed *Cauchy*(0,2), with the exception of α, where selected narrower prior distributions so that the sigmoid-transformed parameters were roughly uniform in [0,1] a priori; prior mean and SD were *Normal*(0,1).

We estimated the joint distribution of the parameters of the model, conditional on all subjects’ observed choices and rewards. For this, we used Markov Chain Monte Carlo (MCMC) techniques (specifically the No-U-Turn variant of Hamiltonian Monte Carlo) as implemented in the Stan modeling language (v2.5, 8). Given a probabilistic generative model (the above equations) and a subset of observed variables, MCMC techniques provide samples from the conditional joint distribution over the remaining latent variables. We ran four chains of 4,000 samples each, discarding the first 2,000 samples of each chain for burn-in. We examined the time-series plots of the chains visually for convergence and also computed Gelman and Rubin’s (9) potential scale reduction factors. For this, large values indicate convergence problems, whereas values near 1 are consistent with convergence. We ensured that these diagnostics were less than 1.02 for all variables.

**Computational Modeling Results for Experiment 2**

Using the complementary computational analysis detailed above, we estimated learning rates and choice stochasticity, in addition to model-based, model-free and exploratory behaviour. This allowed us to test if changes in learning rates and/or choice randomness might explain our findings of increased exploration under CO2. Consistent with the one-trial back regression analysis, CO_2_ had a significant effect on stay/switch behaviour only, such that subjects were more likely to switch to a new action under acute CO_2_ (Table S7). This does not correspond to more randomness in choice, which is captured by the stochasticity parameter.

**Table S6. Group-level estimates of the effect of CO_2_(on/off) on each free parameter in the computational model.**

| **Influence of CO_2_ (ON/OFF) on Model Parameter Estimates** | | | | | |
| --- | --- | --- | --- | --- | --- |
| **Upper 95%**  **Median**  **Lower 95%** | **α**$\mathbf{CO}_{\boldsymbol{2}}$ | ***p***$\mathbf{CO}_{\boldsymbol{2}}$ | $\boldsymbol{mb}\mathbf{CO}_{\boldsymbol{2}}$ | $\boldsymbol{m}\mathbf{fCO}_{\boldsymbol{2}}$ | $\boldsymbol{beta}\boldsymbol{2}\mathbf{CO}_{\boldsymbol{2}}$ |
|  | 0.29 | **-0.04** | 0.07 | 0.23 | 0.12 |
|  | -0.21 | **-0.16** | -0.05 | 0.08 | -0.11 |
|  | -0.60 | **-0.27** | -0.17 | -0.06 | -0.35 |

***α* = learning rate; *p* = perseveration; *mb* = model-based; *mf* = model-free; *beta2* = choice stochasticity.**

**For the effect of CO_2_ on each parameter, the median posterior estimate is given, together with the 95% confidence intervals. Only the slope of *p*CO_2_ (i.e. the effect of CO_2_ on perseveration) is significantly different from zero, such that subjects were more likely to switch choices from trial to trial (i.e. perseverate less) under CO_2_.**

**Experiment 3**

**Exclusion criteria for online task data.** In line with suggestions made for conducting experiments online using Amazon’s Mechanical Turk (AMT), *a priori* exclusion criteria were applied to ensure data quality (10). Subjects were excluded if they missed more than 10% of trials (n=62), responded on the same key on more than 95% of trials on which they registered a response (n=85) or had implausibly fast reaction times, i.e. ±2 standard deviations from the mean (n=18). *Clinical Questionnaires Exclusion Criterion:* In an effort to identify participants who were not reading the questions prior to selecting their responses, we included one catch item: “If you are paying attention to these questions, please select "A little" as your answer”. Very few subjects failed to select the appropriate response to this catch question; those that did were excluded (n=6). *IQ Test Exclusion Criterion:* Participants who did not answer correctly to any of the IQ questions were excluded from further analysis (n=87). The adaptive character of the test meant that participants responding incorrectly received increasingly easy items; consistently failing to respond correctly indicates that given participants might have been inattentive or dishonest. In total, 258/1671 (15%) were excluded from this experiment, in line with a previously published report using this dataset. Note that in this dataset, it was also established that the results did not change regardless of the application of these criteria (11).

**Detailed Results for Model-Based Task and Panic Attacks (past week).** Basic results from this task, and its association to compulsivity, age and IQ, have been published in detail elsewhere (12). The novel results relevant to this study are as follows: one-trial-back regression analysis controlling for IQ, age and gender only, revealed that the frequency of panic attacks in the past week was associated with reductions in model-based planning (p=.012), and also increase in switch behavior (p=.04), but no effect on model-free learning (p=.80). Neither of these significant effects survived inclusion of compulsivity in the model (panic_attack*model-based, *p*=.33; panic_attack*switching, *p*=.24).

**Table S7. Results from Regression Analysis with Anxiety Attacks**

| **Coefficient** | **β** | **SE** | ***z*-value** | ***p*-value** |
| --- | --- | --- | --- | --- |
| **model-based * panic attack** | **-0.03** | **0.01** | **-2.52** | **.012*** |
| *controlling for compulsivity* | -0.01 | 0.01 | -0.97 | .33 |
| model-free * panic attack | -.005 | 0.02 | -0.25 | .80 |
| *controlling for compulsivity* | .004 | 0.02 | 0.223 | .82 |
| **repetition * panic attack** | **-0.07** | **0.04** | **-2.09** | **.04*** |
| *controlling for compulsivity* | -.04 | 0.04 | -1.19 | .23 |

**Detailed Results for Model-Based Task and Life Stress (12 months).**

**Table S8. Results from Regression Analysis with Life Stress (12 month)**

| **Coefficient** | **β** | **SE** | ***z*-value** | ***p*-value** |
| --- | --- | --- | --- | --- |
| **model-based * life stress** | **-.02** | **.01** | **-2.02** | **.04*** |
| *controlling for compulsivity* | -.01 | .01 | -.98 | .33 |
| model-free * life stress | -.01 | .02 | -.74 | .46 |
| *controlling for compulsivity* | -.01 | .02 | -.46 | .65 |
| repetition * life stress | -.02 | .03 | -.87 | .38 |
| *controlling for compulsivity* | -.01 | .03 | -.22 | .83 |

**Computational Modeling Method for Experiment 3**

The computational model proceeded exactly in Experiment 2, except that the within-subject manipulation was absent. We estimated each subject’s learning rate, model-based, model-free, perseveration and choice stochasticity parameters and then tested the extent to which these parameters were associated with panic attacks and life stress, after controlling for age, gender, IQ and the compulsive dimension in secondary regression analyses. The general pattern from the simpler analysis was reproduced with a couple of slight differences. First, the effect of panic attacks on model-based planning was not significant, even without controlling for compulsivity (Table S10). Second, the effect of panic attacks on choice switching (*p*) was significant both when compulsivity was and was-not controlled for (Table S10).

**Table S9. Association between having a recent panic attack (Item 1 on PDSS) and parameters in the computational model.**

| **Coefficient** | **β** | **SE** | ***z*-value** | ***p*-value** |
| --- | --- | --- | --- | --- |
| learning rate * panic attack | .00 | .01 | 0.25 | .81 |
| *controlling for compulsivity* | .01 | .01 | 0.88 | .38 |
| **perseveration * panic attack** | **-0.05** | **0.02** | **-2.55** | **.01**** |
| *controlling for compulsivity* | -0.04 | 0.02 | -1.87 | .06 |
| model-based * panic attack | -0.02 | 0.01 | -1.62 | .10 |
| *controlling for compulsivity* | -0.00 | 0.01 | 0.25 | .81 |
| model-free * panic attack | -0.01 | 0.03 | -0.32 | .75 |
| *controlling for compulsivity* | 0.01 | 0.03 | 0.30 | .76 |
| stochasticity * panic attack | -0.03 | 0.04 | -0.66 | .51 |
| *controlling for compulsivity* | -0.04 | 0.04 | 0.90 | .37 |

**Table S10. Association between Life Stress (12 months) on parameters in the computational model**

| **Coefficient** | **β** | **SE** | ***z*-value** | ***p*-value** |
| --- | --- | --- | --- | --- |
| learning rate * panic attack | -0.00 | .01 | -.02 | .98 |
| *controlling for compulsivity* | 0.00 | .01 | .36 | .72 |
| perseveration * panic attack | -0.00 | .02 | -.18 | .86 |
| *controlling for compulsivity* | 0.01 | .02 | .38 | .70 |
| **model-based * panic attack** | **-0.02** | **.01** | **-2.33** | **.02*** |
| *controlling for compulsivity* | -0.01 | .01 | -1.17 | .24 |
| model-free * panic attack | -0.02 | .02 | -0.61 | .54 |
| *controlling for compulsivity* | -0.01 | .03 | -0.22 | .82 |
| **stochasticity * panic attack** | **-0.08** | **.03** | **-2.43** | **.02*** |
| *controlling for compulsivity* | -0.05 | .03 | -1.52 | .13 |

**References**

1.	Sharp ME, Foerde K, Daw ND, Shohamy D. Dopamine selectively remediates 'model-based' reward learning: a computational approach. Brain. 2016;139(Pt 2):355-64. doi: 10.1093/brain/awv347. PubMed PMID: 26685155.

2.	Daw ND, Gershman SJ, Seymour B, Dayan P, Dolan RJ. Model-Based Influences on Humans' Choices and Striatal Prediction Errors. Neuron. 2011;69(6). doi: 10.1016/j.neuron.2011.02.027. PubMed PMID: WOS:000288886900015.

3.	Rummery G, Niranjan M. On-Line Q-Learning Using Connectionist Systems. Cambridge, UK: Cambridge University Press; 1994.

4.	Lau B, Glimcher PW. Dynamic response-by-response models of matching behavior in rhesus monkeys. J Exp Anal Behav. 2005;84(3):555-79. PubMed PMID: 16596980; PubMed Central PMCID: PMCPMC1389781.

5.	Ito M, Doya K. Validation of decision-making models and analysis of decision variables in the rat basal ganglia. J Neurosci. 2009;29(31):9861-74. doi: 10.1523/JNEUROSCI.6157-08.2009. PubMed PMID: 19657038.

6.	Bellman R. Dynamic Programming. Princeton, NJ: Princeton University Press; 1957.

7.	Otto AR, Raio CM, Chiang A, Phelps EA, Daw ND. Working-memory capacity protects model-based learning from stress. Proc Natl Acad Sci U S A. 2013;110(52):20941-6. doi: 10.1073/pnas.1312011110. PubMed PMID: 24324166; PubMed Central PMCID: PMCPMC3876216.

8.	Stan. Stan: A C++ Library for Probability and Sampling, Version 2.5 available from: <http://mc-stan.org/2014> [October 15, 2014].

9.	Gelman A, Rubin D. Inference from iterative simulation using multiple sequences. Statistical Science. 1992;7(4):457-72.

10.	Crump MJ, McDonnell JV, Gureckis TM. Evaluating Amazon's Mechanical Turk as a tool for experimental behavioral research. PLoS One. 2013;8(3):e57410. doi: 10.1371/journal.pone.0057410. PubMed PMID: 23516406; PubMed Central PMCID: PMCPMC3596391.

11.	Gillan CM, Kosinski M, Whelan R, Phelps EA, Daw ND. Characterizing a psychiatric symptom dimension related to deficits in goal-directed control. Elife. 2016;5. Epub 2016/03/01. doi: 10.7554/eLife.11305. PubMed PMID: 26928075; PubMed Central PMCID: PMCPMC4786435.

12.	Gillan C, Kosinski M, Whelan R, Phelps E, Daw N. Characterizing a psychiatric symptom dimension related to deficits in goal-directed control. eLife. 2016;5(e11305 ). doi: <http://dx.doi.org/10.7554/eLife.11305>.
